# The vitamin K oxidoreductase VKORC1L1 prevents oxidative stress in hepatocytes and protects from MASLD and hepatocellular carcinoma

**DOI:** 10.64898/2026.02.25.707953

**Authors:** Shayesteh Kiani, Julie Lacombe, B. Ashok Reddy, Émilie Gobeil, René P. Michel, Benoit J. Arsenault, Mathieu Ferron

## Abstract

Generation of vitamin K hydroxyquinone (VKH_2_) by vitamin K oxidoreductase 1 (VKORC1) is essential for the γ-carboxylation of clotting factors by hepatocytes. Here, we uncover a non-redundant function of the vitamin K oxidoreductase paralogue VKORC1L1 in liver homeostasis. Mice lacking *Vkorc1l1* globally or specifically in hepatocytes exhibit normal coagulation yet develop progressive metabolic dysfunction–associated steatotic liver disease (MASLD). Transcriptomic profiling revealed early dysregulation of lipid metabolism and inflammatory pathways, converging on human MASLD signatures, while genetic colocalization analyses implicate human *VKORC1L1* variants in MASLD and liver fat accumulation. Mechanistically, VKORC1L1 prevents reactive oxygen species overload and DNA damage through vitamin K reduction, independently of γ-carboxylation. Loss of VKORC1L1 induces oxidative stress, chromosome instability, and aneuploidy, culminating in steatohepatitic hepatocellular carcinoma (HCC). Conversely, pharmacological vitamin K supplementation rescues oxidative stress, MASLD and DNA damage in *Vkorc1l1*-deficient mice. These findings redefine vitamin K as a hepatic antioxidant and identify VKORC1L1 as a safeguard against MASLD and HCC.

## INTRODUCTION

Hepatocellular carcinoma (HCC) is the most common form of primary liver tumour and the third leading cause of mortality by cancer worldwide ^1,2^. Unfortunately, targeted therapies for HCC are limited and prognosis is generally unfavourable as less than 20% of the patients survive more than 5 years following diagnostic. The major causes of HCC are infections with hepatitis viruses (B or C) or exposure to aflatoxin B1, alcohol or other environmental carcinogens ^3^. Another emerging important risk factor for HCC, is metabolic dysfunction-associated steatotic liver disease (MASLD) ^4^. MASLD, defined as an abnormal accumulation of lipids within the liver, can progress to metabolic dysfunction-associated steatohepatitis (MASH), an inflammatory and fibrotic state of the disease which can ultimately evolve to HCC ^5,6^. The mechanisms underlying the progression of MASLD and MASH to HCC remain to be fully uncovered, however it appears that endoplasmic reticulum (ER) stress, inflammation, and oxidative stress might be involved in the induction of DNA damage and hepatocarcinogenesis ^7,8^.

In metazoans, vitamin K (VK) functions in its reduced form (hydroxyquinone; VKH_2_) as a co-factor of the γ-carboxylase (GGCX), an enzyme localized in the ER that converts glutamic acid (Glu) residues to γ-carboxyglutamic acid (Gla) residues in specific proteins transiting through this organelle ^9^. Upon carboxylation of protein, VKH_2_ is oxidized to VK 2,3-epoxide (VKO). VKO is next reconverted to VKH_2_ by the vitamin K oxidoreductase (VKORC1) in a two-step process (i.e., VKO→VK→VKH_2_). Collectively, the actions of GGCX and VKORC1 form the VK cycle. In liver, γ-carboxylation plays a critical role by regulating the function of many coagulation factors (e.g., prothrombin, Factors VII, etc.).

In addition to VKORC1, vertebrates possess a second proteins with VKO reductase activity, a paralogue called VKORC1L1, whose function remains unclear ^10,11^. In vitro and in cell-based assay, both VKORC1 and VKORC1L1 display VKOR activities and support γ-carboxylation of protein ^12,13^. In mice lacking VKORC1, VKORC1L1 can partially support protein carboxylation in the liver during the embryonic and perinatal period. However, beyond postnatal day 7, only VKORC1 can sustain γ-carboxylation and coagulation ^14^. In addition, the sole loss of VKORC1L1 does not affect γ-carboxylation of protein in the liver or coagulation in embryo and newborn ^14^, suggesting that the genuine physiological function of this enzyme remains to be uncovered.

With the objective of revealing the non-redundant role of VKORC1L1 in vivo we herein characterized in details mice lacking this enzyme in all cells, or specifically in hepatocytes. We show that although VKORC1L1 is completely dispensable for γ-carboxylation and coagulation in adult animals, this enzyme is essential to prevent MASLD in mice fed a normal chow diet. This finding was corroborated by the identification of a genetic signal at the human *VKORC1L1* locus colocalizing with MASLD. Our data also establish that through VK reduction, VKORC1L1 protects hepatocytes from reactive oxygen species overload and DNA damage. Consequently, in absence of VKORC1L1, older animals develop HCC tumours which closely recapitulate histological features of human steatotic HCC. Together, this study uncovers a previously uncharacterized role for VKORC1L1 and VK in hepatocytes and establishes that this enzyme has a unique biological function independently of protein γ-carboxylation.

## RESULTS

### *Vkorc1l1* prevent MASLD and MASH without affecting γ-carboxylation

To address the non-redundant function of VKORC1L1 in vivo, we generated *Vkorc1l1^-/-^* mice on a pure C57BL/6J genetic background and carefully monitored their development and general physiology. *Vkorc1l1^-/-^* body weight (BW) was about 20% lower than WT mice at birth and this difference persisted throughout growth and adulthood (Extended Data Fig.1A). A similar difference in BW was previously reported in another loss-of-function model of *Vkorc1l1* in mice and was attributed to reduced adipose tissue mass ^15^. However, in our model, whole-body composition by EchoMRI shows that both fat mass and lean mass were not significantly decreased in *Vkorc1l1*^-/-^ mice when normalized to BW (Extended Data Fig. 1B). In contrast, body length as well as femur and tibia length were significantly reduced in *Vkorc1l1*^-/-^ mice compared to controls (Extended Data Fig. 1C and 1D), suggesting that the absence of VKORC1L1 is causing a general growth defect. At necropsy, all the *Vkorc1l1^-/-^* organs appeared normal and their respective weight was proportional to the body weight, except the liver that was enlarged starting at 6 months of age (Extended Data Fig. 1E). At 12 months of age, the liver of *Vkorc1l1^-/-^* mice was clearly showing sign of MASLD: it was enlarged and pale compared to age-match WT liver (**Fig.1A**, top panels). Histological analyses confirmed the presence of lipid droplets in the liver of these animals (Extended Data Fig. 1F, top panels).

**Figure 1:**
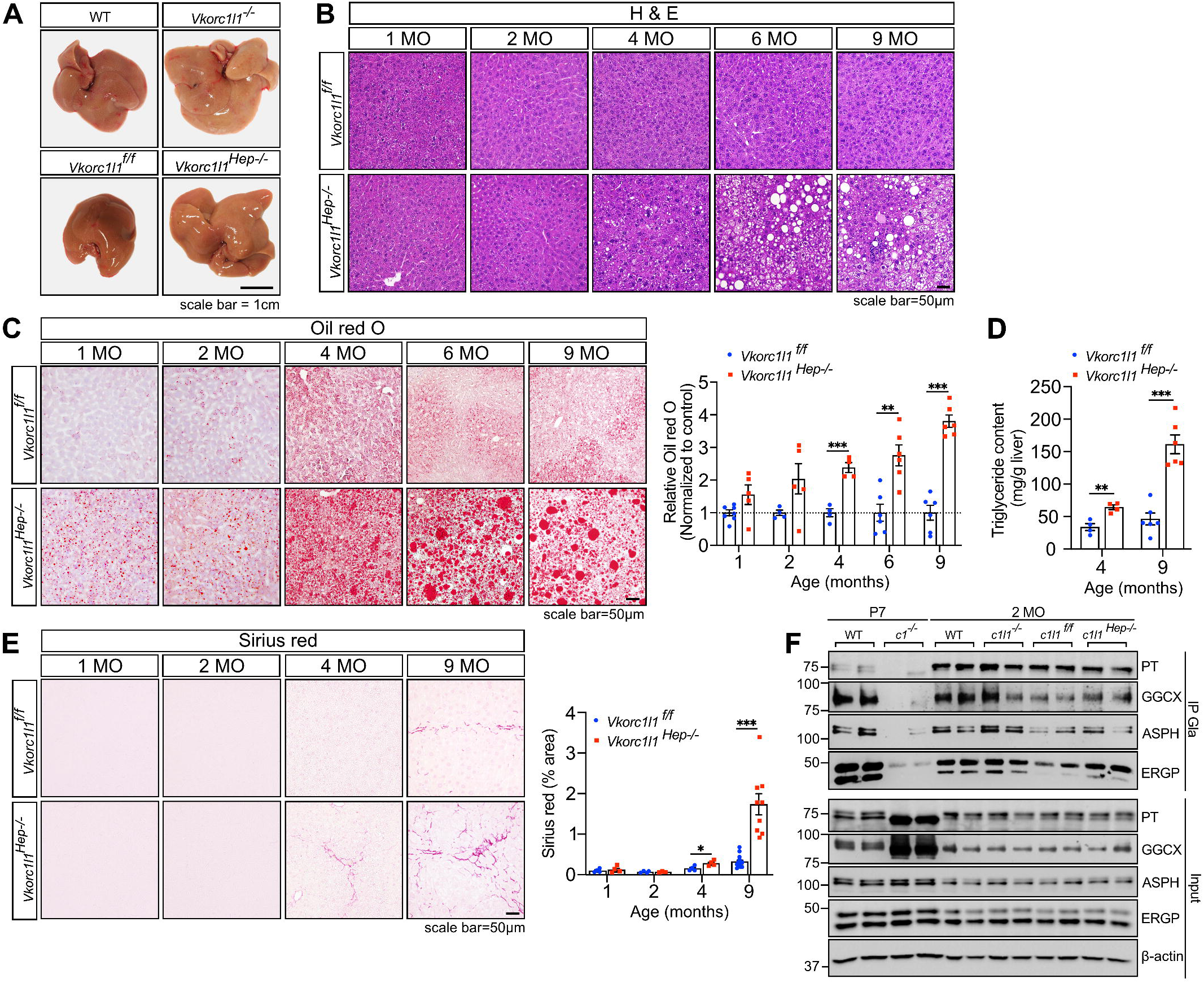
*Vkorc1l1* protects mice from liver steatosis and fibrosis independently of *γ* - carboxylation. **(A)** Gross liver morphology of 1-year-old wildtype (WT), *Vkorc1l1^-/-^, Vkorc1l1^f/f^* and *Vkorc1l1 ^Hep-/-^* male mice. **(B)** Hematoxylin and eosin (H&E) staining of liver sections from *Vkorc1l1^f/f^* and *Vkorc1l1 ^Hep-/-^* male mice of the indicated ages. **(C)** Oil red O staining of liver cryosection from *Vkorc1l1^f/f^* and *Vkorc1l1 ^Hep-/-^* male mice of the indicated ages (left). The quantifications (n = 4-7) are presented as a fold change relative to *Vkorc1l1^f/f^* control (right). **(D)** Liver triglyceride content of 4- and 9-month-old *Vkorc1l1^f/f^* and *Vkorc1l1 ^Hep-/-^* male mice. Content was normalized to the tissue weight (n = 4-6). **(E)** Sirius red staining of the liver sections from *Vkorc1l1^f/f^* and *Vkorc1l1 ^Hep-/-^* male mice of the indicated ages (left). The quantifications (n = 4-12) are presented as the percentage of Sirius red area over total liver area (right). **(F)** Prothrombin (PT), GGCX, ASPH and ERGP γ -carboxylation in liver extract from WT and *Vkorc1^-/-^* (*c1^-/-^*) postnatal day 7 (P7) pups, and from WT, *Vkorc1l1^-/-^* (*c1l1^-/-^*)*, Vkorc1l1^f/f^* (*c1l1^f/f^*) and *Vkorc1l1 ^Hep-/-^* (*c1l1 ^Hep-/-^*) 2-month-old mice was assessed by immunoprecipitation with anti-Gla antibody followed by Western blot analyses. Inputs represent 2% of total protein extracts. All mice were maintained on a regular chow diet. Results represent the mean ± SEM. Multiple unpaired, 2-tailed Student’s t test was used in **(C)**, **(D)**, and **(E)**. ***p < 0.001, **p < 0.01, *p < 0.05.

Based on these observations, we next generated mice lacking *Vkorcl1* specifically in hepatocytes (*Vkorc1l1^Hep-/-^* mice) and assessed the impact on body size and development of MASLD. *Vkorc1l1^flox/flox^* mice ^16^ were crossed with *Alb-Cre* mice expressing the Cre recombinase under the control of the albumin promoter ^17^ (Extended Data Fig. 1G and 1H). The BW of *Vkorc1l1^Hep-/-^* mice did not differ from their control littermates (i.e., *Vkorc1l1^flox/flox^*) at any of the ages examined (Extended Data Fig. 1I). However, the weight of their liver was increased at 6 months of age with signs of steatosis (**Fig.1A**, bottom panels, and Extended Data Fig. 1J). The presence of abnormal lipid droplets in the liver of *Vkorc1l1^Hep-/-^* mice was detected histologically using hematoxylin and eosin (H&E) and Oil Red O staining as early as 2 months of age and gradually increased as the mice aged (**Fig.1B-C**). Accordingly, liver triglycerides content was increased by about 2 and 3-fold at 4 and 9 months respectively (**Fig. 1D**). *Vkorc1l1^+/+^;Alb-Cre* mice had normal liver weight compared to wildtype (*Vkorc1l1^+/+^*) or *Vkorc1l1^flox/flox^* mice, and no sign of steatosis was detected histologically in these animals at 9 months of age (Extended Data Fig. 1K), ruling out the possibility that the *Alb-Cre* transgene itself was responsible for the observed phenotype. Glucose tolerance, insulin sensitivity and gluconeogenesis were not affected in *Vkorc1l1^Hep-/-^* mice at 4 months of age (Extended Data Fig. 1L-N), suggesting that hepatic insulin resistance was not involved in the observed phenotype. Hence, inactivation of *Vkorc1l1* in hepatocytes triggers MASLD without impacting body growth, body weight and insulin sensitivity, suggesting that VKORC1L1 regulates hepatocytes function through a cell-autonomous mechanism.

The progression of MASLD to MASH in absence of VKORC1L1 was next evaluated thrpugh Sirius red staining. This revealed the presence of mild fibrosis at 4-month of age in *Vkorc1l1^Hep-/-^* mice, while more extensive fibrosis was detected at 9 months in the same mice (**Fig.1E**). Fibrosis was also present in the liver of older *Vkorc1l1*^-/-^ mice (Extended Data Fig. 1F, bottom panels).

VK reduction in hepatocytes is essential for the γ-carboxylation and activity of several clotting factors, including prothrombin, and this process can be partially supported by VKORC1L1 during development and perinatally in the absence of VKORC1 ^14^. We therefore evaluated the impact of *Vkorc1l1* inactivation on coagulation and hepatic γ-carboxylation in adult mice. Adult *Vkorc1l1*^-/-^ mice have normal prothrombin time as assessed by the measurement of the international normalized ratio (Extended Data Fig. 1O). An assay involving immunoprecipitation of carboxylated proteins using rabbit α-Gla antibody, followed by western blot analysis, revealed that γ-carboxylation of prothrombin and GGCX occurs normally in the liver of adult *Vkorc1l1*^-/-^and in *Vkorc1l1^Hep-/-^* mice, but was greatly reduced in 1 week old *Vkorc1*^-/-^ pups as expected (**Fig.1F**). Similar results were obtained when assessing the γ-carboxylation of aspartate-beta-hydroxylase (ASPH) and Endoplasmic Reticulum Gla Protein (ERGP), two recently identified intracellular VK-dependent proteins ^18^. Together, these biochemical analyses suggest that *Vkorc1l1* deficiency causes MASLD without affecting VK-dependent carboxylation in hepatocytes.

### Loss of *Vkorc1l1* in hepatocytes triggers transcriptional changes associated with MASLD and MASH in humans

To better characterized the metabolic changes associated with the inactivation of *Vkorc1l1* in hepatocytes, we analyzed by RNA-sequencing the transcriptomic profile of *Vkorc1l1^Hep-/-^* liver at 1, 2 and 9 months of age. These different time points were selected to detect transcriptional alterations preceding the appearance of MASLD and MASH, i.e., 1 and 2 months of age, or associated with the advanced stages of the phenotype, i.e., 9 months of age. As expected from the gradual appearance of the steatosis and fibrosis, the number of differentially expressed genes (DEG; adjusted *P* value ≤ 0.05) increased markedly from 155 at 1 month, to 694 at 2 months and 1972 at 9 months (Supplementary Tables 1-3). Gene set enrichment analysis revealed that several biological processes (BP) linked to lipid metabolism and fatty acid metabolism, lipid biosynthesis, biosynthesis of unsaturated fatty acids, and carboxylic acid metabolism were significantly enriched in the DEG at all ages (**Fig.2A**), suggesting dysregulated fatty acid synthesis and transport in absence of VKORC1L1 in hepatocytes. Processes associated with the cellular response to xenobiotic metabolism were also dysregulated in 1 month old *Vkorc1l1^Hep-/-^* liver. Several of the DEG associated with xenobiotic metabolism encoded for cytochromes P450 (*Cyp2d9*, *Cyp2b9*, *Cyp2b10*, etc.) previously associated with the metabolism of fatty acids.

**Figure 2:**
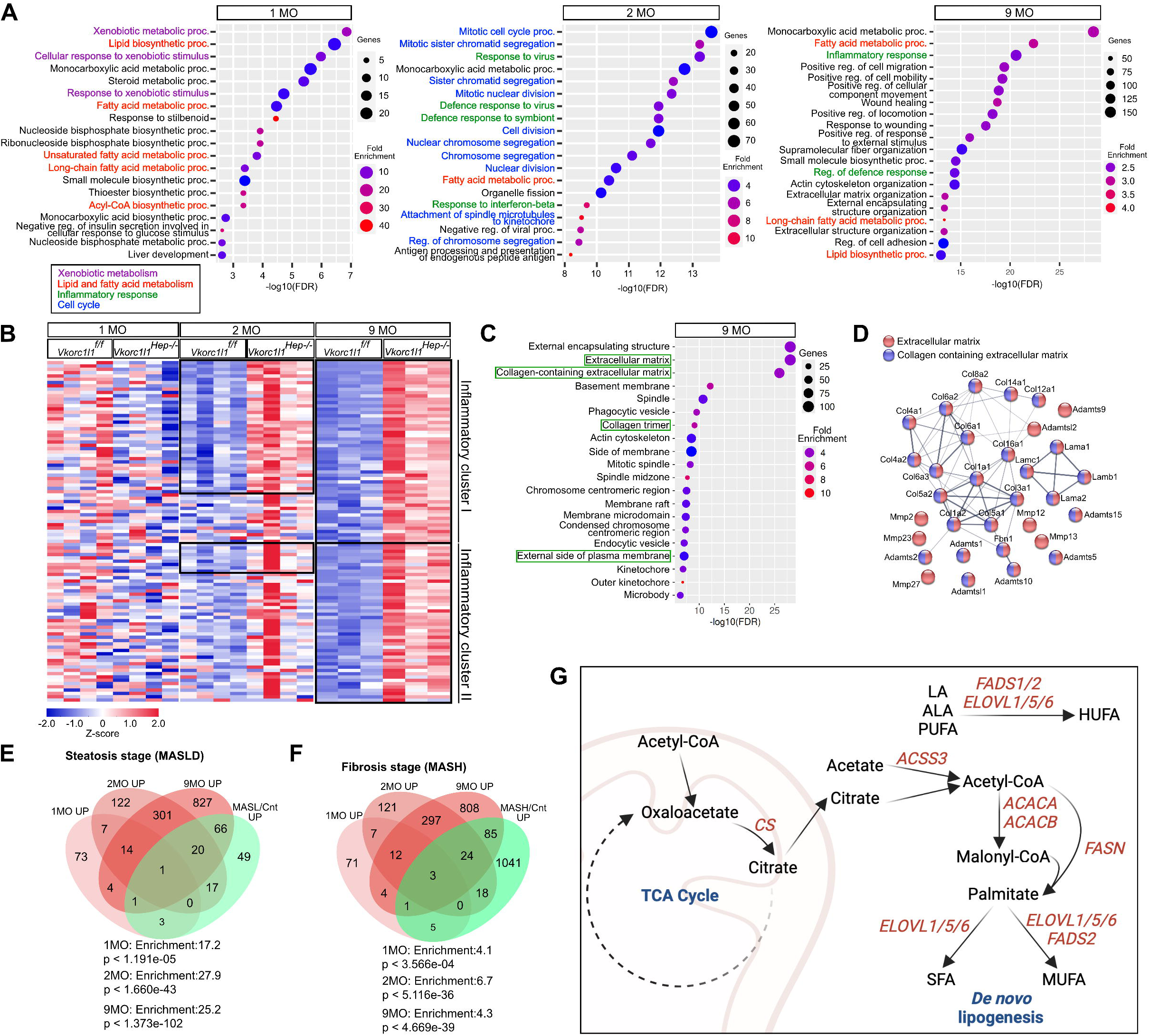
Absence of *Vkorc1l1* in hepatocytes is associated with steatosis, fibrosis and inflammation gene signatures. Gene set enrichment analysis was performed for differentially expressed genes (DEGs) in *Vkorc1l1 ^Hep-/-^* compared to *Vkorc1l1^f/f^* male mice using Gene ontology (GO). **(A)** Dot plots representing GO analysis of DEGs in *Vkorc1l1 ^Hep-/-^* livers at 1, 2 and 9 months of age using biological processes terms (BP). **(B)** Heatmaps of up-regulated inflammatory clusters at 1, 2 and 9 months of age. **(C)** Dotplot representing GO analysis of up-regulated genes in *Vkorc1l1 ^Hep-/-^* liver at 9 months of age using cellular component terms (CC). **(D)** Gene networks of up-regulated genes in 9-month-old *Vkorc1l1 ^Hep-/-^* liver associated with extracellular matrix and collagen. **(E-F)** Venn diagrams showing the overlap between up-regulated genes in 1-, 2- and 9-month-old *Vkorc1l1* deficient livers and up-regulated genes in human MASLD **(E)** and MASH **(F)**. Number of genes is shown for each overlap. **(G)** Schematic representation of de novo lipogenesis pathway. The genes highlighted in red are up-regulated in both human MASLD and *Vkorc1l1 ^Hep-/-^* liver. **(A)** and **(C)**, data are presented as -log_10_(FDR) and number of genes and fold enrichment are indicated with bullet size and color respectively.

There was an enrichment at 2- and 9-month-old in *Vkorc1l1^Hep-/-^* liver for biological processes linked to immune or inflammatory process, such as response to virus, response to interferon-beta and inflammatory response (**Fig. 2A**). Further unsupervised clustering analyses focusing on the up-regulated genes at 2 and 9 months, revealed two clusters of up-regulated inflammatory genes (cluster I with 39 and 55 genes; cluster II with 9 and 48 genes at 2 and 9 months respectively). The genes in these clusters are associated with interferon and cytokine signaling (e.g., *Irf9*, *Eif2ak2*, *Ifngr1*, *Il10rb*, etc.) and chemokine signaling pathway (e.g. *Ccr5*, *Ccl22*, *Cxcl16*, etc.) respectively, suggesting progressive immune cell infiltration and inflammation in the *Vkorc1l1^Hep-/-^* liver (**Fig.2B**; Supplementary Table 4), a feature associated with MASH in human^19^.

In the group of DEG at 9 months there was a strong enrichment for biological processes such as wound healing, supramolecular fiber organization and extracellular matrix organization (**Fig.2A**). We observed that up-regulated genes at 9 months of age were strongly enriched for cellular components (GO:CC) of the extracellular matrix, collagen-containing extracellular matrix and extracellular space (**Fig. 2C**). These gene signatures included genes encoding for several collagens (*Col1a1*, *Col1a2*, *Col3a1*, *Col6a1, Col6a2*, etc.), matrix metalloproteinases (*Mmp2, Mmp13, Mmp13*, etc.), other metalloproteinases of the ADAMTS family (*Adamts1, 2, 5, 10, 15,* etc.) and additional extracellular matrix proteins such as fibrillin (*Fbn1*) and laminins (*Lama1, Lama2, Lamb1, Lamc1*), suggesting extensive remodeling of the extracellular matrix in livers in absence of VKORC1L1 (**Fig. 2D**). These transcriptional changes are also consistent with the presence of fibrosis in *Vkorc1l1^Hep-/-^* liver at this age (**Fig.1E**).

We next determined to which extent the transcriptome of *Vkorc1l1^Hep-/-^* liver intersects with the gene expression profile of human MASLD and MASH. These analyses revealed a progressive and significant overlap between the genes dysregulated at 1, 2 and 9 months of age in *Vkorc1l1^Hep-/-^* liver and a previously established gene signature in human MASLD and MASH ^20^ (**Fig.2E-F;** Extended Data Fig. 2A & B). Interestingly, the expression of *VKORC1L1* was reduced in average by 35% in human MASH (adjusted *P* value = 1.02x10^-06^), but not in MASLD, compared to the control healthy subjects (Extended Data Fig. 2B). Several of the genes up-regulated in both human MASLD/MASH and *Vkorc1l1^Hep-/-^* mice encode for proteins involved in fatty acid synthesis [acetyl-CoA carboxylase 1 and 2 (*Acaca* and *Acacb*), acyl-CoA synthetase short-chain family member 3 (*Acss3*), and fatty acid-synthase (*Fasn*)], elongation [elongation of very long chain fatty acids protein 1, 5 and 6 (*Elovl1*, *Elovl5* and *Elovl6*)] or desaturation [fatty acid desaturase 1 and 2 (*Fads1* and *Fads2*) (**Fig.2G**). Therefore, the absence of VKORC1L1 trigger transcriptional changes that recapitulate human MASLD and MASH.

### A genetic variant in human *VKORC1L1* is associated with liver fat in humans

In parallel, we investigated potential genetic evidence linking *VKORC1L1* to MASLD in humans. A large genetic association study identified a genetic signal for MASLD at the *GUSB* locus, which lies in the vicinity of *VKORC1L1* ^21^. Mendelian randomization (MR) and genetic colocalization analyses combining an eQTL dataset obtained from 504 liver samples ^22^ along with an updated MASLD GWAS meta-analysis ^23^ prioritized *VKORC1L1* over *GUSB* as the potential causal gene for both MASLD and liver fat proton density fat fraction (PDFF) (Table 1). For MASLD, *VKORC1L1* (top SNP rs12666915-T) showed a significant MR association (β_MR_ = –0.199, P_MR_ = 0.004) without evidence of pleiotropy (P_HEIDI_= 0.228), whereas *GUSB* (rs75312426-C) did not reach significance (P_MR_ = 0.434). Similarly, for liver fat measured by PDFF, *VKORC1L1* (rs12666915-T) was significantly associated (β_MR_= –0.131, P_MR_ = 0.005) with no pleiotropy (P_HEIDI_ = 0.018), while *GUSB* showed no significant effect (P_MR_ = 0.195). Colocalization analyses further supported *VKORC1L1*, with posterior probabilities of shared causal variants (PP.H4) of 35.4% for MASLD and 34.4% for liver fat, compared to <2% for *GUSB*. Together, these findings implicate *VKORC1L1*, rather than *GUSB*, as the more likely effector gene at this locus for MASLD and liver fat accumulation.

**Table 1.**
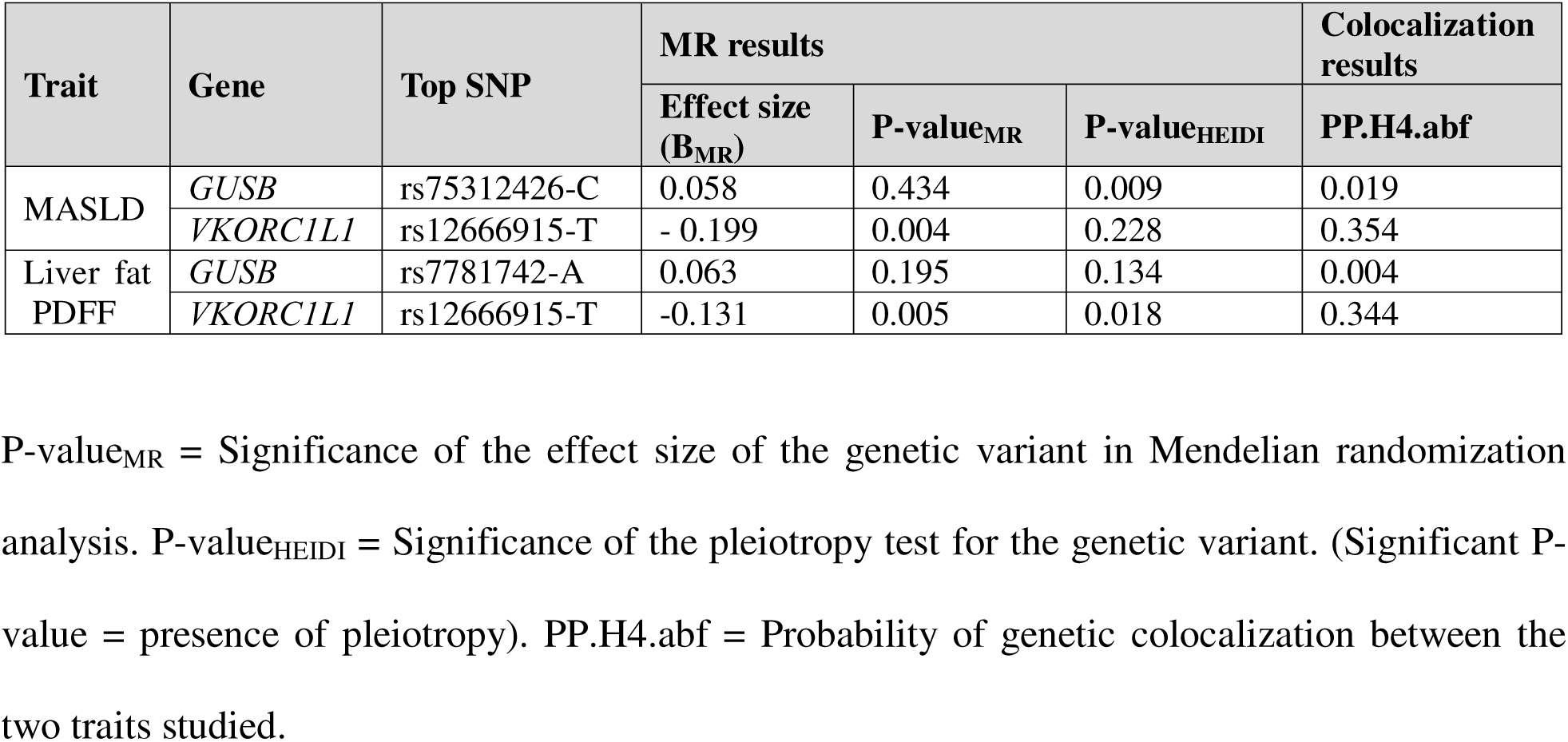
Mendelian randomization and genetic colocalization prioritize *VKORC1L1* over *GUSB* at the 7q11.21 locus for MASLD and liver fat.

| Trait | Gene | Top SNP | MR results |  |  | Colocalization results<br>PP.H4.abf |
| --- | --- | --- | --- | --- | --- | --- |
| | | | Effect size ($B_{MR}$) | P-value<sub>MR</sub> | P-value<sub>HEIDI</sub> | |
| MASLD | *GUSB* | rs75312426-C | 0.058 | 0.434 | 0.009 | 0.019 |
|  | *VKORC1L1* | rs12666915-T | - 0.199 | 0.004 | 0.228 | 0.354 |
| Liver fat PDFF | *GUSB* | rs7781742-A | 0.063 | 0.195 | 0.134 | 0.004 |
|  | *VKORC1L1* | rs12666915-T | -0.131 | 0.005 | 0.018 | 0.344 |

P-value<sub>MR</sub> = Significance of the effect size of the genetic variant in Mendelian randomization analysis. P-value<sub>HEIDI</sub> = Significance of the pleiotropy test for the genetic variant. (Significant P-value = presence of pleiotropy). PP.H4.abf = Probability of genetic colocalization between the two traits studied.

### VKORC1L1 inactivation is associated with a chromosome instability signature and signs of aneuploidy

Gene ontology analysis of the transcriptomic data at 2 MO of age also indicates that *Vkorc1l1^Hep-/-^* livers are characterized by the upregulation of a large gene network comprising genes previously involved in mitotic cell cycle process, cell division, mitotic spindle assembly, DNA repair and the regulation of the G2/M transition of the cell cycle (**Fig.3A**). This gene cluster included centromere proteins (*Cenpc1, Cenpe, Cenpf, Cenpi*, etc.), kinesin-like protein involved in spindle formation or cytokinesis (*Kif15, Kif20a*, *Kif20b, Kif22, etc.*), component of the kinetochore (*Nuf2, Ndc80, Spc24, etc.*) and mitotic checkpoint regulators (*Aurkb, Bub1, Bub1b, Ccnb1, Ccnb2, Mad2l1, Ttk, etc.*). Several genes (e.g., *Foxm1*, *Ttk*, *Kif20a*, *Rad51ap1*, etc.) in this network are part of a chromosomal instability (CIN) gene signature previously associated with aneuploidy, tumorigenesis and/or poor clinical outcome in multiple human cancers, including HCC (**Fig.3B-C**) ^24,25^.

**Figure 3:**
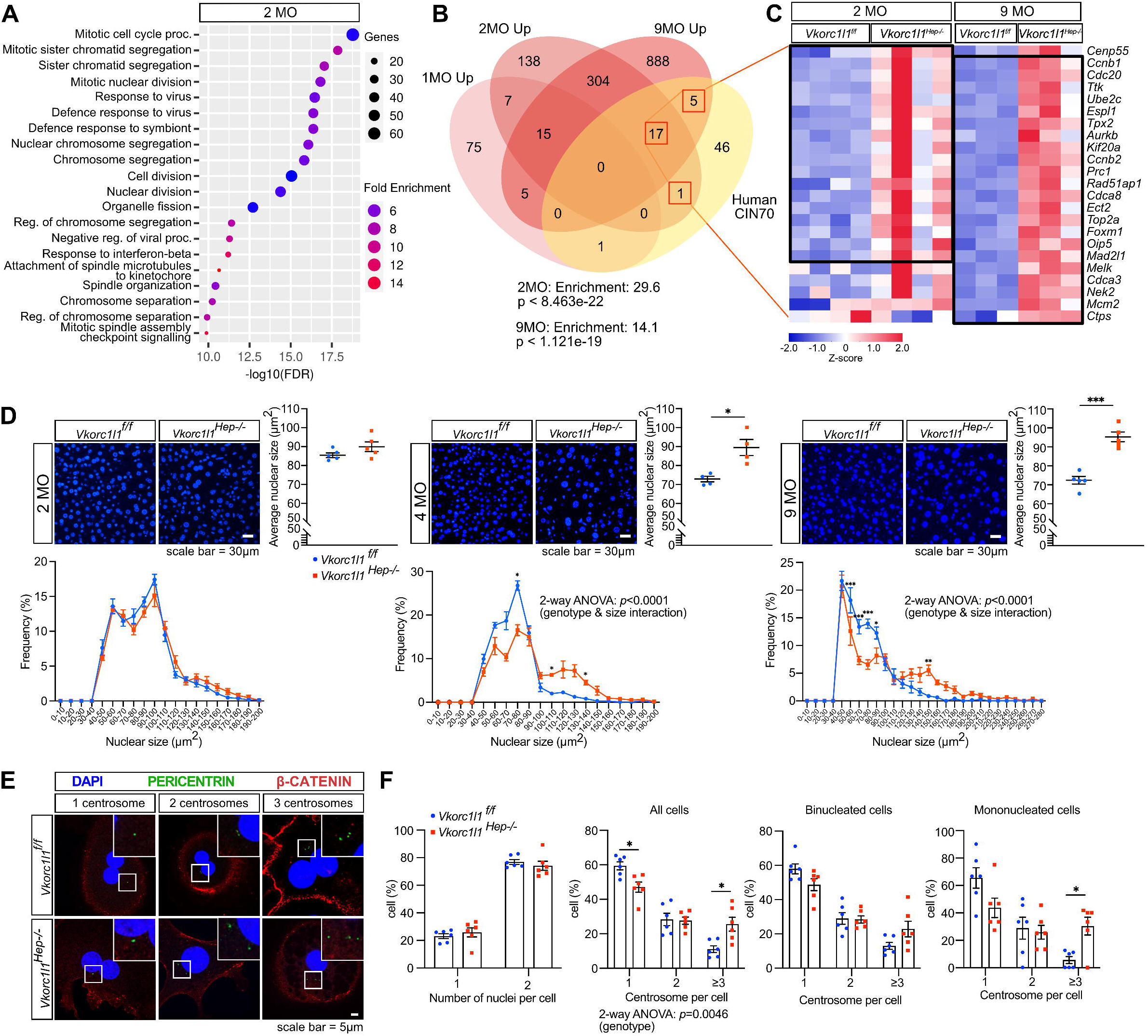
Inactivation of *Vkorc1l1* in hepatocytes is associated with signs of chromosome instability. **(A)** Dot plot representing Gene Ontology (GO) analysis of up-regulated genes in 2-month-old *Vkorc1l1 ^Hep-/-^* male livers using biological processes terms (BP). Circle sizes represent the number of genes associated with the GO, and the color represent the fold enrichment. **(B)** Overlap between a previously described human chromosome instability signature (CIN70) ^24^ and the up-regulated genes in 1-, 2- and 9-month-old *Vkorc1l1 ^Hep-/-^* male livers. **(C)** Heatmap of the up-regulated genes both in CIN70 and in 2- and 9-month-old *Vkorc1l1 ^Hep-/-^* male livers. **(D)** Representative DAPI staining of the nuclei on liver sections from 2-, 4- and 9-month-old *Vkorc1l1^f/f^* and *Vkorc1l1 ^Hep-/-^* male mice. The average nuclear size and the distribution of nuclei size are represented (n = 4-5). **(E)** Immunofluorescence staining for the centrosome marker pericentrin on primary hepatocytes isolated from *Vkorc1l1^f/f^* and *Vkorc1l1 ^Hep-/-^* male mice (n = 6). Nuclei and centrosomes number per cell in hepatocytes from *Vkorc1l1^f/f^* and *Vkorc1l1 ^Hep^*^-/-^ mice were quantified (n=242-263). Results represent the mean ± SEM. Unpaired, 2-tailed Student’s t test was used in **(D)**. Two-way ANOVA with Bonferroni’s post tests was used in **(D)** and in **(E)**. ***p < 0.001, **p < 0.01, *p < 0.05.

Histological examination of liver sections revealed increased proportion of larger nuclei and more heterogeneous nuclear size in *Vkorc1l1^Liv-/-^* hepatocytes compared to controls at 4 and 9 months of age (**Fig.3D**). These results suggest increased nuclear ploidy or karyomegaly in absence of VKORC1L1 and are coherent with nuclear enlargement in human MASLD ^26,27^. Centrosome amplification which is often associated with CIN ^28^ was analyzed in hepatocytes isolated from 4-month-old mice by immunofluorescence of pericentrin, a centrosome specific protein. As expected, based on previous studies, about 75% of control hepatocytes were binucleated, and this proportion was not changed in the *Vkorc1l1^Hep-/-^* hepatocytes (**Fig.3E**). The number of cells with 1 centrosome was reduced, while the number of cells with 2 centrosomes remained unchanged in the *Vkorc1l1^Hep-/-^* cells compared to control. However, the percentage of cells with 3 or more centrosomes was significantly increased in the *Vkorc1l1^Hep-/-^* hepatocytes population. Moreover, this increase in the number of cells with ≥3 centrosomes was observed specifically in the mononucleated hepatocytes subpopulation, supporting the presence of abnormal multipolar spindle formation and CIN in a subset of *Vkorc1l1^Hep-/-^* hepatocytes (**Fig.3E-F**). A persistent population of Ki-67 positive hepatocytes in *Vkorc1l1^Hep-/-^* liver after 2 months of age also suggest anomalies in cell cycle progression (Extended Data Fig. 3A). Altogether, these results indicate that VKORC1L1 prevents abnormal cell cycle progression and aneuploidy in hepatocytes.

### Global or liver specific inactivation of *Vkorc1l1* causes HCC

MASH can progress to HCC in humans and VKORC1L1 inactivation in hepatocytes was associated with molecular and cellular signs of genomic instability. We therefore next investigated if aging *Vkorc1l1^-/-^* and *Vkorc1l1^Hep-/-^* mice develop liver cancer. Several *Vkorc1l1^-/-^* mice started to show swollen abdomen around 18 months, and a few died before 24 months. Necropsy revealed the presence of liver tumours in 18- to 24-month-old (MO) *Vkorc1l1^-/-^* and *Vkorc1l1^Hep-/-^* mice (**Fig.4A**). This phenotype was fully penetrant in males as almost all the 18-24 MO *Vkorc1l1^-/-^* and *Vkorc1l1^Hep-/-^* mice examined (n=11-24 mice per genotype for each strain) harbored several macroscopic tumours in their liver (**Fig.4B**). No tumor was found in WT littermates and only two *Vkorc1l1^f/f^* littermates had a single tumor. Histopathological analysis revealed that these tumors were of grade 1 to 3, with a majority of grade 2 in both models (**Fig.4C**), and were HCC of the “steatohepatitic variant” (**Fig.4D left panels,** Extended Data Fig. 3B), a type of HCC frequently associated with MASLD and MASH ^29,30^. Additional histological analyses, revealed high proliferation index (Ki67) (**Fig.4D right panels**), and elevated expression of markers of human HCC such as α-fetoprotein (AFP), GRP78 and cytokeratin 19 (CK19) ^31–34^ (**Fig.4E**, Extended Data Fig. 3B).

**Figure 4:**
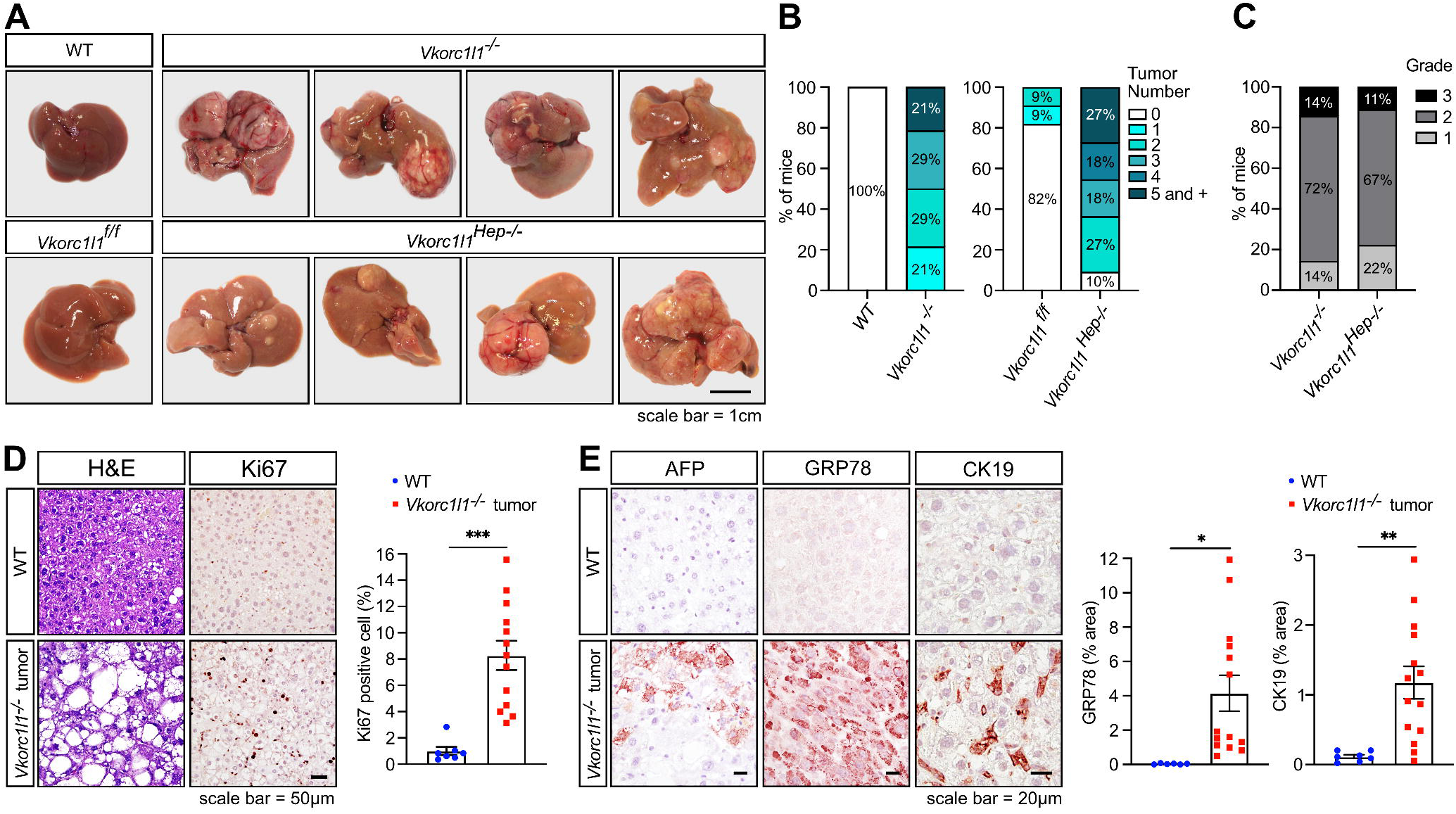
Global and liver-specific inactivation of *Vkorc1l1* induce hepatocellular carcinoma in mice. **(A)** Gross liver morphology of 18- to 24-month-old *Vkorc1l1^-/-^* and *Vkorc1l1 ^Hep-/-^* male mice, and WT and *Vkorc1l1^f/f^* control littermates. **(B)** Tumor frequency and number in *Vkorc1l1^-/-^* and *Vkorc1l1 ^Hep-/-^* male mice. **(C)** Histologic grading of HCC tumors from *Vkorc1l1^-/-^* and *Vkorc1l1 ^Hep-/-^* male mice from 1 to 3 (1: well-differentiated; 2: moderately differentiated; 3: poorly differentiated). **(D)** H&E staining (left) and Ki67 immunohistochemistry (right) on liver or tumor sections from 18- to 24-month-old WT and *Vkorc1l1^-/-^* male mice. Quantification represents percentage of Ki67 positive cells over total cell number (n = 7-13). **(E)** Alpha fetoprotein (AFP), glucose-regulated protein 78 (GRP78) and Cytokeratin 19 (CK19) immunohistochemistry on liver or tumor sections from 18- to 24-month-old WT and *Vkorc1l1^-/-^* male mice. Quantifications are shown as the percentage of GRP78 or CK19 area over total liver area (n = 6-14). Results represent the mean ± SEM. Unpaired, 2-tailed Student’s t test was used in **(D)** and **(E)**. ***p < 0.001, **p < 0.01, *p < 0.05.

### VKORC1L1 protects hepatocytes from oxidative stress and DNA damage

To understand mechanistically how VKORC1L1 may prevent MAFLD and tumorigenesis in hepatocytes, we examined transcriptomic data at 1 month of age, before the appearance of steatosis or CIN. At this age, only 155 genes are dysregulated, however, a strong enrichment for genes encoding proteins with oxidoreductase activity was detected (Extended Data Fig. 4A). This suggested that the absence of VKORC1L1, which itself acts as an oxidoreductase, resulted in dysregulated oxidoreduction processes and oxidative stress in hepatocytes. Since VKH_2_ has recently emerged as a potent radical trapping antioxidant (RTA) ^35^, we hypothesized that VKORC1L1 may be involved in an antioxidant pathway in hepatocytes through the generation of VKH_2_. Supporting this notion, a previously described hepatocytes oxidative stress gene signature^36^ was progressively induced in *Vkorc1l1^Hep-/-^* liver (**Fig.5A**). Moreover, 1-month-old *Vkorc1l1^Hep-/-^* isolated hepatocytes were characterized by an increased level of intracellular reactive oxygen species (ROS) measured with a fluorogenic probe (CellROX Green; **Fig.5B**; Extended Data Fig. 4B). The same cells were also more sensitive to the induction of ROS by treatments with a low dose of *tert*-Butyl hydroperoxide (t-BHP) ^37^, suggesting that VKORC1L1 protects hepatocytes from an exogenous inducer of oxidative stress (**Fig.5B**; Extended Data Fig. 4B). The presence of oxidative stress was next assessed in vivo in *Vkorc1l1^Hep-/-^* liver using an antibody recognizing 8-hydroxy-2’deoxyganosine (8-OHdG) and 8-hydroxyguanosine (8-OHG), which directly reflect oxidative damage to DNA and RNA in cells ^38^. These immunofluorescence staining indicated high 8-OHdG/8-OHG levels as early as 2 weeks of age in *Vkorc1l1^Hep-/-^* liver and a progressive increase as the animals aged (**Fig.5C**).

**Figure 5:**
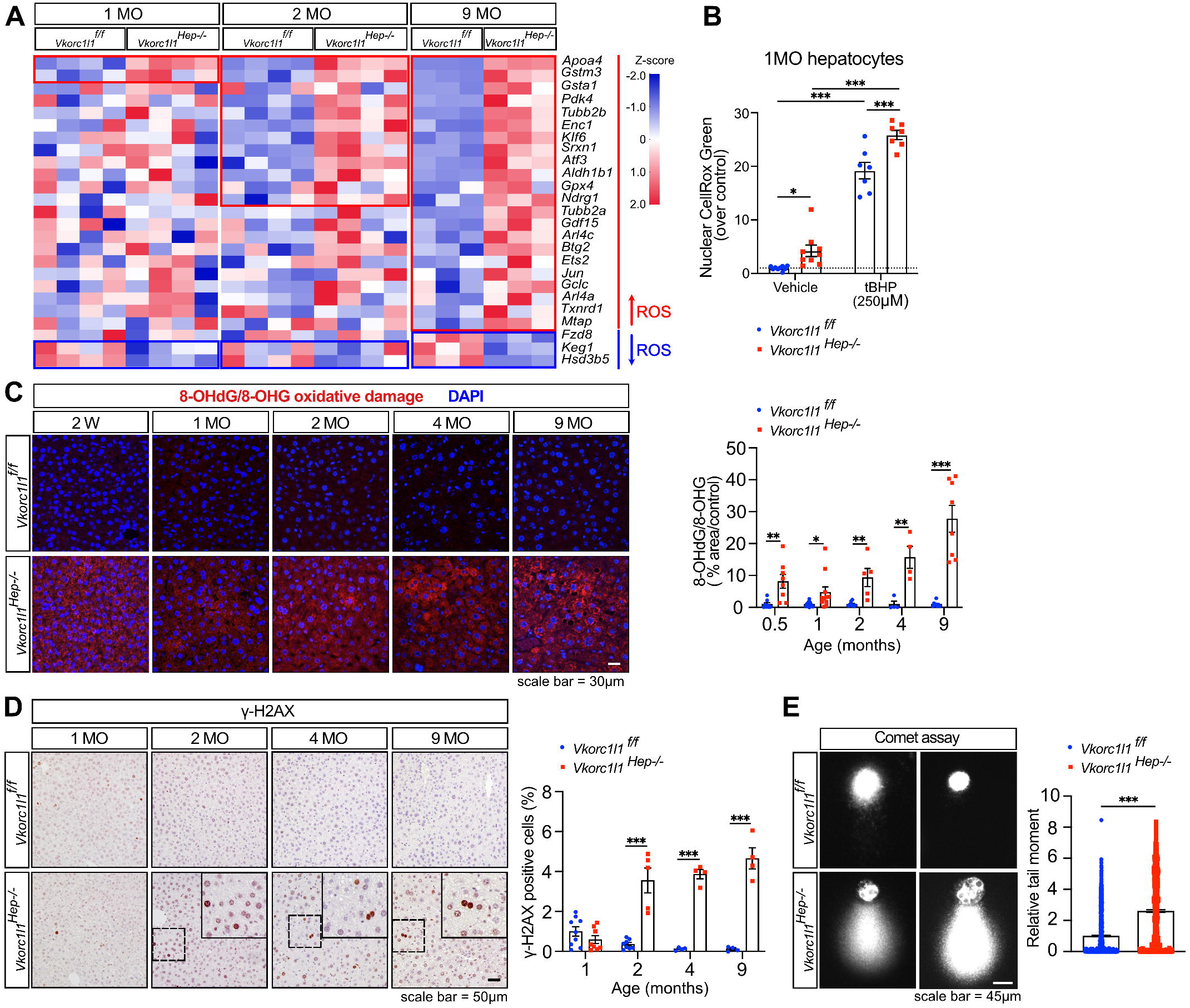
*Vkorc1l1*-deficient mice showed increased level of oxidative stress, and DNA damage. **(A)** Heatmap representation of a previously described hepatocyte oxidative stress gene signature ^36^ in *Vkorc1l1^f/f^* and *Vkorc1l1 ^Hep-/-^* male mice livers at indicated ages (n =3-4). Red indicates significantly up-regulated genes, and blue significantly down-regulated genes. **(B)** ROS measurement using the CellROX Green fluorometric probe in primary hepatocytes isolated from 1-month-old *Vkorc1l1^f/f^* and *Vkorc1l1 ^Hep-/-^* male mice, treated or not with 250 μM *tert*-butyl hydroperoxide (tBHP) for 2 hours. CellROX Green signal in DAPI+ nuclei was quantified and is presented as a fold change relative to *Vkorc1l1^f/f^* control without tBHP treatment (n=7-9 experimental replicates per condition from 4 independent experiments). **(C)** Immunofluorescence staining of 8-hydroxy-2’deoxyguanosine (8-OHdG) and 8-hydroxyguanosine (8-OHG), an oxidative stress marker, on liver sections from *Vkorc1l1^f/f^* and *Vkorc1l1 ^Hep-/-^* male mice at the indicated ages. Quantifications are presented as the percentage of 8-OHdG/8-OHG signal over total liver area and normalized by *Vkorc1l1^f/f^* at each age (n = 4-8). **(D)** γ-H2AX immunohistochemistry on liver sections from *Vkorc1l1^f/f^* and *Vkorc1l1 ^Hep-/-^* male mice at the indicated ages. Quantification represent percentage of γ-H2AX positive cells over total cell number (n = 4-9). **(E)** Comet assay on primary hepatocytes isolated from 4-month-old *Vkorc1l1^f/f^* and *Vkorc1l1 ^Hep-/-^* male mice (n = 3-5 mice; >750 comets per genotype were quantified). Results represent the mean ± SEM. Two-way ANOVA with Fisher’s LSD post-tests was used in **(B)**. Unpaired, 2-tailed Student’s t test were used in **(C-E)**. ***p < 0.001, **p < 0.01, *p < 0.05.

Elevated cellular ROS-induced single-strand DNA lesions such as 8-OHdG can produce double-strand break (DSB) in genomic DNA, potentially leading to oncogenic events ^39^. Accordingly, phosphorylated histone H2AX (γ-H2AX), a histological marker of DSB, was detected at high level in the nucleus of *Vkorc1l1^Hep-/-^* hepatocytes in vivo starting at 2 months of age (**Fig.5D**). The presence of DSB in *Vkorc1l1^Hep-/-^* liver was confirmed directly using “comet” assays on isolated 4-month-old hepatocytes (**Fig.5E**). Together, these data establish that VKORC1L1 plays an essential role in protecting hepatocytes from ROS accumulation and DNA damage in vivo.

A study suggesting that VKORC1L1 may protect cancer cells from ferroptosis, a non-apoptotic form of cell death involving lipid peroxidation ^40^ prompted us to examinate if this pathway could be driving MASLD in *Vkorc1l1^Hep-/-^* mice. However, levels of malondialdehyde (MDA), a byproduct of lipid peroxidation, were unaffected in *Vkorc1l1^Hep-/-^* livers compared to controls at all ages tested (Extended Data Fig. 4C). Lipid peroxidation was also assessed directly in primary hepatocytes using a sensitive fluorescent probe, BODIPY 581/591 C11 undecanoic acid. This revealed that control and *Vkorc1l1^Hep-/-^* hepatocytes had the same levels of lipid peroxidation whether they were treated with vehicle or with RSL3, a pharmacological activator of ferroptosis (Extended Data Fig. 4D). Therefore, although VKORC1L1 protect hepatocytes from oxidative stress, it does not appear to prevent ferroptosis in the same cells.

### Exogenous vitamin K rescues the absence of VKORC1L1 ex vivo

The bleeding defects associated with warfarin overdosing or *VKORC1* inactivation in mice and humans can be rescued by treatment with large doses of vitamin K_1_ or K_2_ ^11,41,42^. This was recently explained by the enzymatic action of FSP1 (encoded by *AIFM2*), a warfarin-resistant NAD(P)H-dependent vitamin K quinone reductase also expressed in hepatocytes ^35,43^. Importantly, although FSP1 can catalyze the reduction of VK to VKH_2_, it is unable to convert VKO to VK (**Fig.6A**) ^35^. We therefore tested the capacity of VKO and VK to reverse oxidative stress in primary 4-month-old *Vkorc1l1*-deficient hepatocytes. The results indicate that ROS are more elevated at baseline in *Vkorc1l1^Hep-/-^* primary hepatocytes as observed in 1-month-old hepatocytes (**Fig. 5B** and **6B**). Interestingly, both phylloquinone (VK_1_) and menaquinone (VK_2_) significantly reduced ROS levels in *Vkorc1l1^Hep-/-^* hepatocytes when provided at a high concentration (20μM), suggesting the presence of another VK reductase with lower activity in the cells which could be either VKORC1 or FSP1. In contrast, VK_1_O was ineffective at reducing ROS in *Vkorc1l1^Hep-/-^* hepatocytes, demonstrating that VKORC1 is unable to compensate for the lack of VKORC1L1 activity. Importantly, VK_1_, VK_2_ and VK_1_O equally increased γ-carboxylation in control and *Vkorc1l1^Hep-/-^* hepatocytes, indicating normal GGCX and VKORC1 function in absence of VKORC1L1 (Extended Data Fig. 5A). Together, these data indicate that VKORC1L1 is the main VK epoxide reductase protecting hepatocytes from oxidative stress and that exogenous VK can partially rescue the absence of VKORC1L1 in vitro.

**Figure 6:**
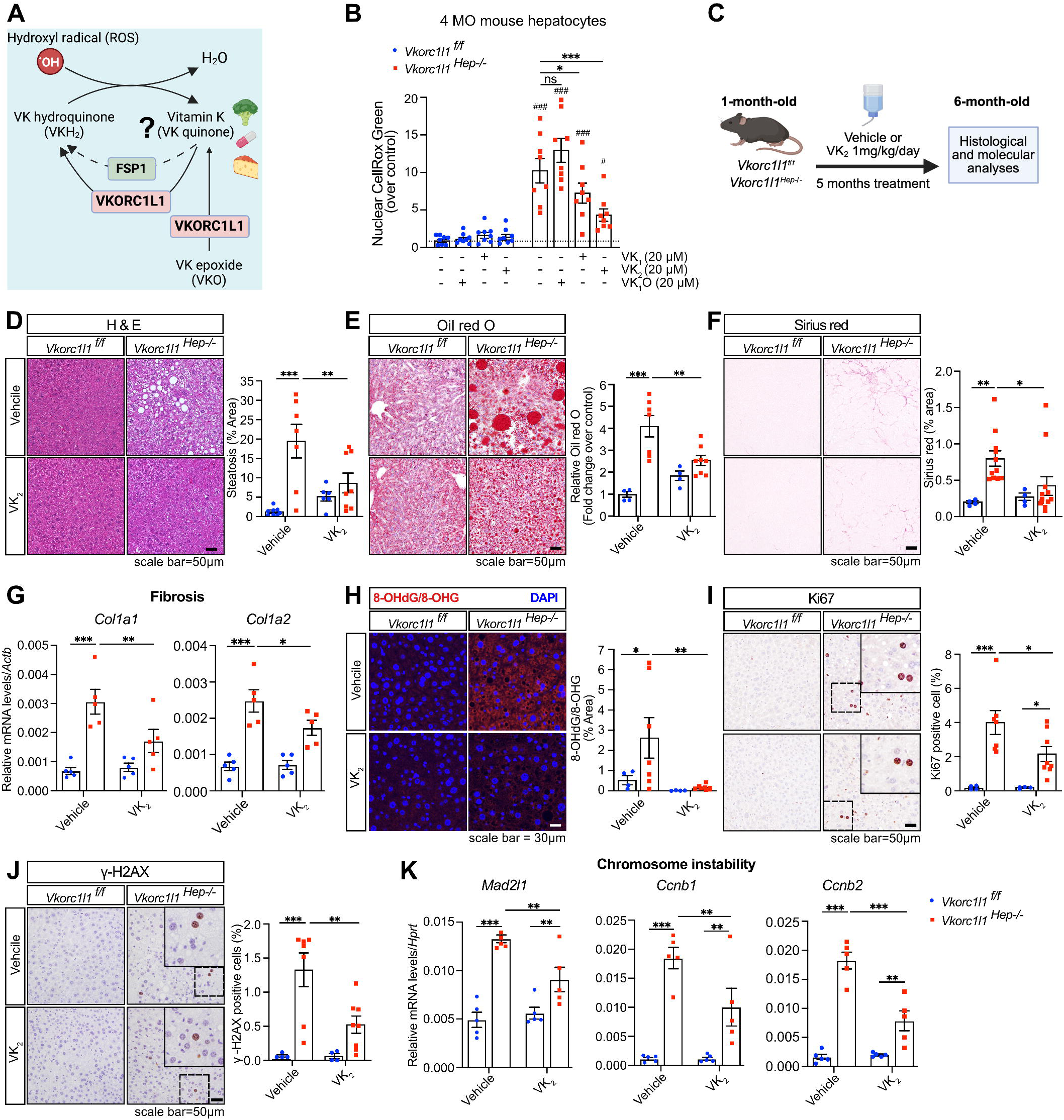
Vitamin K rescues oxidative stress in *Vkorc1l1* deficient hepatocytes and prevent steatosis, fibrosis and oxidative stress in *Vkorc1l1 ^Hep-/-^* mice. **(A)** Model depicting the role of VKORC1L1 and FSP1 as mediator of the antioxidant function of vitamin K in hepatocytes. **(B)** Hepatocytes isolated from 4-month-old *Vkorc1l1^f/f^* and *Vkorc1l1 ^Hep-/-^* male mice were cultured for 24h with 20 µM of phylloquinone (VK_1_), menaquinone-4 (VK_2_) or VK_1_ 2,3-epoxide (VK_1_O) and ROS measured using CellROX green fluorometric probe. Signal in DAPI+ nuclei was quantified and normalized to *Vkorc1l1^f/f^* untreated hepatocytes (n=8 experimental replicates per condition from 3 independent experiments). (C-K) *Vkorc1l1^f/f^* and *Vkorc1l1 ^Hep-/-^* male mice fed a chow diet were treated with vehicle or VK_2_ in drinking water (1mg/kg/day)) for 5 months starting at 1 month of age **(C)**. **(D)** H&E staining of liver section (n = 6-8). **(E)** Oil red O staining of liver cryosections (n = 4-8). **(F)** Sirius red staining of liver sections (n = 4-8). **(G)** Gene expression analysis of fibrotic genes by qPCR on the liver mRNA (n = 5). **(H)** Immunofluorescent staining on liver sections of 8-hydroxy-2’deoxyganosine (8-OHdG) and 8-hydroxyguanosine (8-OHG) (n=4-8). **(I)** Ki67 immunohistochemistry on the liver sections (n = 4-8). **(J)** γ-H2AX immunohistochemistry on liver sections (n=4-8). **(K)** Gene expression analysis of chromosome instability genes by qPCR on liver mRNA (n = 5). Results represent the mean ± SEM. Two-way ANOVA with Fisher’s LSD was used in **(B, D-K)**. In **(B)** ###p < 0.001, ##p < 0.01, #p < 0.05 when comparing *Vkorc1l1^f/f^* and *Vkorc1l1 ^Hep-/-^* hepatocytes in the same experimental condition. ***p < 0.001, **p < 0.01, * p < 0.05.

### Pharmacological doses of VK rescue VKORC1L1 deficiency in vivo

We next tested if the phenotypes observed in *Vkorc1l1^Hep-/-^* mice could be prevented by treatment with high doses of VK in vivo. Since both VK_1_ and VK_2_ could dampen oxidative stress in *Vkorc1l1^Hep-/-^* hepatocytes we first examined the bioavailability of these two forms when provided in the drinking water. A dose of 9 mg/L (20μM), equivalent to 1 mg/kg/day considering a mouse drink approximately 4mL of water per day, was initially administered to control mice for 2 weeks and levels of γ-carboxylated GGCX and ERGP in the liver assessed by western blotting as a proxy for VK bioavailability in this organ. Since this analysis revealed that VK_2_ was more efficient at increasing γ-carboxylation in the liver (Extended Data Fig. 5B), 1-month-old control or *Vkorc1l1^Hep-/-^* mice were provided with VK_2_ or vehicle in drinking water continuously for 5 months (**Fig.6C**). The treatment had no impact on body weight (Extended Data Fig. 5C) but significantly increased γ-carboxylation in the liver in both controls and *Vkorc1l1^Hep-/-^* mice (Extended Data Fig. 5D). At the end of the study, histological and gene expression analyses were performed to assess steatosis, fibrosis, oxidative stress and DNA damage.

Steatosis and fibrosis which were significantly increased in vehicle treated *Vkorc1l1^Hep-/-^* mice, were normalized to control levels by VK_2_ treatment (**Fig.6D-F;** Extended Data Fig. 5E). Quantitative PCR shows that the expression of *Col1a1* and *Col1a2* which is increased in *Vkorc1l1^Hep-/-^* liver and associated with MASH in humans ^44^, was reduced significantly in the *Vkorc1l1^Hep-/-^* animals treated with VK_2_ (**Fig.6G**). VK_2_ completely rescued the DNA and RNA oxidative damage, assessed by 8-OHdG/8-OHG staining, in *Vkorc1l1^Hep-/-^* liver (**Fig.6H**). In addition, the percentage of γ-H2AX and Ki67 positive cells was reduced by about 50% in the livers of *Vkorc1l1^Hep-/-^* mice receiving VK_2_ compared to vehicle treated animals (**Fig.6I-J**). Expression of the chromosome instability markers *Mad2l1*, *Ccnb1* and *Ccnb2* which was increased in *Vkorc1l1^Hep-/-^* livers, was also significantly reduced upon VK_2_ treatment (**Fig.6K**). Altogether, these data indicate that a high dose of exogenous VK_2_ can partially or completely prevent most of the molecular and cellular defects associated with the absence of VKORC1L1. These results also support the conclusion that VKORC1L1 antioxidant function is mediated by VK reduction.

## DISCUSSION

### Vitamin K, MASLD and HCC

A few studies previously suggested the existence of a pathophysiological link between VK cycle, MASLD and hepatocellular carcinoma (HCC). Firstly, ∼35 years ago it was discovered that des-γ-carboxy prothrombin, the undercarboxylated form of prothrombin, is abundantly secreted by HCC cells in culture and is abnormally elevated in the plasma of patients with HCC, suggesting an intrinsic defect in VK metabolism in HCC ^45–49^. Secondly, VK content is reduced in HCC tumours compared to surrounding non-cancerous livers ^50,51^. Thirdly, in the context of a small clinical trial, VK_2_ supplementation reduced significantly the incidence of HCC in women with viral liver cirrhosis ^52^. Fourthly, recent epidemiological studies suggest that inadequate VK intake increases the risk of MASLD in humans ^53,54^. Although these studies suggest that a dysfunction of the VK cycle may be linked to MASLD and HCC, whether the VK cycle directly impacts these diseases was not established. Moreover, the mechanism by which VK may protect hepatocytes from MASLD and HCC was unknown.

Here, using two different mouse models, combined with a large genetic association study, we identify VKORC1L1 as a mediator of the protective effect of VK on MASLD and HCC. *Vkorc1l1^Hep-/-^* mice phenotype recapitulate several features of human MASLD and MASH, including progressive steatosis and fibrosis. In addition, transcriptomic data suggests the presence of inflammation following the first signs of steatosis. Of note, *Vkorc1l1^Hep-/-^* mice develop MASLD and HCC in absence of obesity or diabetes, and without exogenous carcinogen (e.g., diethylnitrosamine). Therefore, at least in mice, VKORC1L1 plays a non-redundant and essential role in protecting hepatocytes from steatosis and carcinogenesis in physiological conditions.

A significant overlap between the transcriptome of *Vkorc1l1^Hep-/-^* liver and human MASLD or MASH suggest that the molecular and cellular mechanisms implicated in this mouse model are similar to the ones driving the human disease. That *VKORC1L1* expression was significantly reduced in human MASH compared to healthy liver, indicate that VKORC1L1 may be implicated in the progression of MASLD to MASH in humans. Moreover, we revealed genetic colocalization between liver expression of *VKORC1L1* and MASLD in large human cohorts. As mentioned above, human HCC is often characterized by an intrinsic defect in VK cycle which is revealed by the presence of des-γ-carboxy prothrombin in the circulation ^45–49^. Yet, mutations in *VKORC1L1* are relatively rare in HCC tumors. It is therefore possible that a defect in VK metabolism or import in hepatocytes contribute to steatosis and carcinogenesis in a significant fraction of human HCC.

### Antioxidant function of VK and VKORC1L1 in normal and cancer cells

In addition to its key function in supporting protein γ-carboxylation, VKH_2_ has recently emerged as an important radical trapping antioxidant (RTA) molecule acting as an inhibitor of ferroptosis, a form of necrotic cell death characterized by oxidative damage to phospholipids. FSP1, encoded by the *AIFM2* gene, was identified as an NAD(P)H-dependant VK reductase able to generate VKH_2_ which can trap lipid peroxyl radicals mediating lipid peroxidation in melanoma cells and in liver ^35^. Additional work has established that FSP1 is the warfarin resistant enzyme responsible for the generation of VKH_2_ from VK in the absence of VKORC1 in HEK 293 cells and likely in vivo in liver ^43^. Another study suggested that VKORC1L1 may protect some cancer cell lines from ferroptosis by generating VKH_2_ ^40^. Importantly, in these different studies, the anti-ferroptotic effect of VK, FSP1 or VKORC1L1 could only be revealed when ferroptosis was induced by genetic or pharmacological inhibition of glutathione peroxidase 4 (GPX4), a master regulator of this cell death pathway.

Our own data indicate that VKORC1L1 protect hepatocytes from ROS by reducing VK and VKO, to generate VKH_2_. These finding are in line with earlier work demonstrating an antioxidant effect of VKH_2_ in vitro ^55,56^. We also show that MDA level was identical in the liver of control and *Vkorc1l1^Hep-/-^* mice and that RSL3, a pharmacological inhibitor of GPX4, equally induced lipid peroxidation in control or VKORC1L1-deficient hepatocytes. These data suggest that VKORC1L1 does not play an essential role in ferroptosis in hepatocytes in physiological conditions. Therefore, depending on cell type and context, VKORC1L1 and VKH_2_ may be protecting from carcinogenesis by preventing DNA damage due to ROS or protecting cancer cells from cell death by ferroptosis. Such a differential and apparently conflicting role on cancer progression and survival for an antioxidant pathway is not unprecedented. Nuclear factor erythroid 2 (Nrf2), a transcription factor controlling the expression of an antioxidant genetic program, has beneficial effects in normal and healthy liver by preventing ROS, steatosis and fibrosis ^57,58^. However, *NFR2* amplification or gain-of-function mutations are frequently observed in human HCC tumors and could drive the oncogenic process by protecting the cancer cells from excessive oxidative stress ^59^.

### Why vertebrates have two VKORCs?

VKOR homologues are present in most metazoans, but also in several protists, bacteria and higher plant species, indicating that VKORs are evolutionary very ancient. γ-carboxylase homologues are only present in metazoans, suggesting that VK reduction as first evolved in prokaryote to fulfil a function different from protein carboxylation. In bacteria, VKORs homologues are implicated in the formation of protein disulfide bond formation in the periplasm^60^. It is interesting to note that plant VKORs homologues have been shown to protect the organism from ROS induced by osmotic stress ^61^. The presence of two VKOR paralogues, VKORC1 and VKORC1L1, is unique to vertebrate genomes and is probably the result of a gene duplication event that occurred in the primitive vertebrate after divergence from the urochordate lineage. Interestingly, sequences of vertebrate VKORC1L1 orthologues are significantly more conserved than the sequences of vertebrate VKORC1 orthologues ^10^. These phylogenetic studies suggest that following gene duplication, VKORC1 was freer to diverge than VKORC1L1. One possible hypothesis to explain these observations is that the housekeeping function of the ancestral VKOR was retained by VKORC1L1, while VKORC1 has diverged to acquire a new function in supporting protein carboxylation. We believe our findings support this hypothesis for several reason. First, VKORC1L1 inactivation in mice does not impact γ-carboxylation in the liver, except in embryo and only if VKORC1 is also inactivated ^14^. Second, mice lacking VKORC1L1 in all cells or only in hepatocytes are characterized by MASLD and HCC, even though VKORC1 and γ-carboxylation are still present in the same cells. Third, VKORC1 deletion is sufficient to block γ-carboxylation in the liver and in osteoblast ^14,16^. Therefore, even if VKORC1 and VKORC1L1 share the same enzymatic activity and are both expressed in hepatocytes, they clearly fulfill different cellular functions.

How two proteins with the same enzymatic activity can perform such different cellular functions remains unknown at this point. Although both VKORC1 and VKORC1L1 are described as ER resident proteins, they may be localized in different microdomains in the ER membrane or be associated with distinct protein complexes within those membranes. That VKORC1 and VKORC1L1 maintain separated pools of VKH_2_ for different cellular function is an attracting hypothesis since VKH_2_ is a hydrophobic membrane bound compound with a short half-life and a limited capacity to diffuse away from its site of production. Future studies aiming at characterizing VKORC1L1 unique protein interacting partners should provide insights on the mechanism by which it prevents ROS in cells.

In conclusion, we propose here that VKORC1L1 prevent steatosis and carcinogenesis in the liver by reducing oxidative stress through the generation of VKH_2_. We also show that this in vivo function of VKORC1L1 occurs independently of γ-carboxylation and cannot be fulfilled by other VK reductases in physiological conditions. Together, our findings extend the metabolic functions of VK and reveal how it may prevent MASLD and HCC.

## Supporting information

Extented data figures 1-5

## ACKNOWLEDGEMENTS

We thank A. Germain for INR measurement, R. Essalmani for technical help with hepatocyte isolations, J. Estall for providing the *Alb-Cre* mice, V. Calderon for RNA-Seq analysis and G. Sanguineti and V. Cornish for providing vitamin K epoxide. We also thank the staff of IRCM Microscopy, Molecular Biology and Histology Core Facilities for their technical support. We acknowledge the invaluable collaboration of the surgery team, bariatric surgeons, and staff of the Quebec Obesity Biobank and would like to thank all study participants from the Institut universitaire de cardiologie et de pneumologie de Québec (IUCPQ).This work was supported by funding from the Cancer Research Society (MF), the Canadian Institutes of Health Research (MF, PJT-195967 and DCP-192065), and the CMDO Network (MF). SK received scholarships from IRCM, McGill University and the Fonds de recherche du Québec – Santé (FRQS). E.G, holds a Doctoral Research Award from the Canadian Institutes of Health Research. B.J.A. holds a Senior Scholar Award from the FRQS.

## Author contributions

J.L. and M.F. conceived the study, designed the experiments, and initiated the project. S.K., J.L., E.G., A.B.R. and M.F. collected and analyzed data. R.P.M. graded the tumors on histological sections. M.F. wrote the manuscript, with the support of S.K. and B.J.A., and all authors commented and contributed to editing the final version. M.F. acts as the guarantor of this work and is responsible for data access.

## METHODS

### Mice

*Vkorc1l1^-/-^* and *Vkorc1l1^f/f^* mice were previously described ^16^. To generate *Vkorc1l1* hepatocyte specific knock out (*Vkorc1l1^Hep-/-^*^)^, *Vkorc1l1^f/f^* mice were crossed with the *Alb-cre* line ^17^. All mice were maintained on a C57BL/6J genetic background, kept in a pathogen-free facility at IRCM on a 12h light/dark cycle and fed a normal chow diet (Teklad gobal 18% protein rodent diet; 2918; Envigo). In some experiments, mice were supplemented with vehicle, VK_1_ (V3501, Millipore Sigma) or VK_2_ (V9378, Millipore Sigma) in the drinking water (∼1 mg/kg/day) for 2 weeks or 5 months starting at weaning. All animal use complied with the guidelines of the Canadian Committee for Animal Protection and was approved by IRCM institutional animal care committee.

### Metabolic tests

Glucose tolerance test (GTT) and pyruvate tolerance test (PTT) were performed after 5 hours of fasting. Blood glucose was measured at baseline and 15-, 30-, 60- and 120-minutes following i.p. injection of glucose (2g/kg of BW) or pyruvate (2g/kg of BW). Insulin tolerance tests (ITT) were performed after 5 hours of fasting. Blood glucose was measured at baseline and 30-, 60-, 90- and 120-minutes following i.p. injection of recombinant insulin (Humulin R; Eli Lilly) (0.75U/kg).

### EchoMRI body composition analysis

Body composition of mice was assessed using an EchoMRI body composition analyzer (Minispec LF-50, Bruker). Mice were placed in a plastic cylinder insert without anesthesia. Total body fat mass, lean mass, free water, and total body water were measured non-invasively according to the manufacturer’s instructions.

### Liver histology

A piece of liver tissue was fixed in 10% formalin for 24 hours at room temperature, embedded in paraffin and sectioned at 5μm. For immunohistochemistry and immunofluorescence experiments, rehydration was followed by an antigen retrieval step (sub-boiling for 10 minutes in 10mM sodium citrate pH 6.0). For immunohistochemistry, antibodies were detected using Vectastain Elite ABC-peroxidase kit (Vector Laboratories; PK-6101) and NovaRED Substrate Kit (Vector Laboratories; SK-4800) following manufacturer’s instructions. For immunofluorescence, blocking was performed in PBS containing 5% normal donkey serum and 0.3% Triton for 1 hour at room temperature. Sections were then incubated with primary antibodies diluted in PBS, 1% BSA and 0.1% Triton for 1 hour at room temperature or overnight at 4°C, and secondary Cy3 donkey anti-mouse (715-165-150; Jackson Immunoresearch Laboratories) and Alexa-Fluor 488-conjugated donkey anti-rabbit (711-545-152; Jackson Immunoresearch Laboratories) antibodies for 1 hour at room temperature. Nuclei were stained with DAPI. The antibodies used in immunohistochemistry and immunofluorescence staining are listed in Supplementary Table 5. Hematoxylin and eosin (H&E), Sirius Red, Masson’s trichrome and Oil Red O stainings were carried out using standard protocols.

Liver cryoblocks were prepared by fixation in 10 ml ice-cold 4% paraformaldehyde for 4 hours at 4 °C, washing three times in PBS (5 minutes each) and incubating in 30% sucrose in PBS overnight at 4 °C. Liver pieces were embedded and frozen in Tissue-Plus^TM^ OCT compound (23730571; Fisher scientific) using dry ice and stored at -80 °C. Tissues were sectioned at 12 μm and stained for Oil Red O.

Quantification of the DNA/RNA oxidative damage, hematoxylin and eosin (H&E), Sirius red and Oil Red O staining was done using Image J. Ki67 and γ-H2AX quantification was performed using OsteoMeasure Analysis System. Volocity 6.0 was used to measure nuclei sizes of DAPI stained liver sections.

### Hepatocyte isolation

Mice were anesthetized by intraperitoneal (i.p.) injection of a single dose of drug mixture of ketamine hydrochloride and xylazine. The liver was perfused from the vena cava with 20 mL of buffer containing 0.2 g/L EGTA, 140 mM NaCl, 7 mM KCl, 10 mM HEPES and 1M NAOH to chelate calcium and wash out blood. Extracellular matrix was dissociated using 15 mL of digestion buffer (140 mM NaCl, 7 mM KCl, 1M NAOH, 4.76 mM CaCl_2_ and 10 mM HEPES) containing 1.5 mg Liberase^TM^ TL (Roche Applied Science). The liver was harvested, ruptured and cells released into low glucose DMEM (5.5 mM glucose). The cells suspension was passed through a 70 μm filter and centrifuged at 50 × *g* for 2 minutes (brake and accelerator set at zero throughout the experiment). Cells were resuspended in 10 ml of low glucose DMEM and 10 mL of Percoll solution (9 volumes of Percoll + 1 volume of 10X PBS) (P4937; MilliporeSigma) was added to the cells. Cells were centrifuge at 200 × *g* for 10 minutes, washed once with 20 mL of low glucose DMEM and counted before platting.

### ROS assay

Glass coverslips (12mm diameter, #1.5 thickness; NEU-GG-12-1.5-OZ; Mandel) were coated with 0.1mg/mL poly-L lysine (P1274; MilliporeSigma) in sterile ddH_2_O at room temperature for 1 hour, washed three times with sterile ddH_2_O, left to dry for 1 hour and 60 000 hepatocytes were plated on each coverslips. Three hours after plating, DMEM was replaced by William’s E with or without vitamin K, and the next day, when required, 250 µM tert-Butyl hydroperoxide (tBHP) was added for 2 hours at 37 °C. Then, media was removed and cells loaded with 5 µM CellROX^TM^ Green probe (C10444, Thermo Fisher) in DMEM for 30 minutes. The cells were washed once with PBS, fixed in 4% paraformaldehyde in PBS for 15 minutes at room and washed twice with PBS. Nuclei were stained with DAPI and the coverslips were mounted on microscope slides with FluorSave reagent (345789; MilliporeSigma).

Imaging was performed using Zen v3.4 on a confocal rotary disk inverted microscope from Zeiss equipped with a Yokogawa CSU-1 module using a 20× objective. Cells not loaded with the probe were used as a control to set threshold below the non-specific signal. Upon oxidation, the CellROX^TM^ Green probe localizes to the nucleus; therefore, the nuclear signal area was quantified. In brief, Volocity 6.0 quantification module was used to threshold for the CellROX^TM^ Green signal and the area of its intersection with the DAPI stained nuclear region was measured. The total CellROX^TM^ Green signal area within the nucleus was normalized to the total nuclear area. Two to three coverslips were quantified per condition in three separate experiments including three mice from each genotype.

### Comet assay

Twenty thousand freshly isolated hepatocytes were mixed with 120 µl of 0.5% low melting agarose and placed on a glass slide coated with 1.5% normal melting agarose. After overnight incubation in lysis solution (1% triton, 10% DMSO, 2.5 M NaCl, 100 mM EDTA, 10 mM Tris, pH 10.0), the slides were placed in alkaline buffer (0.4 M Tris-HCl, pH 7.5) for 1 hour to allow unwinding of DNA. Electrophoresis was conducted at 25V and 300A for 20 minutes in an ice bath at 4 °C, using electrophoresis buffer containing 300mM NaOH and 1mM EDTA. Slides were coated with drops of neutralizing buffer (0.4 M Tris–HCl, pH 7.5) 3 times for 5 minutes each, rinsed with distilled water and air dried at room temperature. Slides were rinsed briefly with distilled water before adding 30 µl of ethidium bromide solution (20µg/ml), then covered with coverslips and imaged using a fluorescence microscope ^62^. The relative tail moment of individual comets was quantified using OpenComet software ^63^.

### Triglyceride content measurement

100mg of liver tissue was washed with cold PBS, homogenized in 1 ml of 5% NP-40 solution using Dounce homogenizer and the assay was performed according to manufacturer’s protocol (ab65336; Abcam).

### Lipid peroxidation measurement

Around 1-10mg of liver tissue was washed with cold PBS, homogenized with using Dounce homogenizer and 3,4-methylenedioxyamphetamine (MDA) level was measured according to manufacturer’s protocol (ab118970, Abcam). All values were normalized to the protein content measured using Bradford protein assay.

Lipid peroxidation was measured in primary hepatocytes using the ratiometric fluorometric probe BODIPY^TM^ 581/591 C11 (D3861, Thermo Fisher). The assay was performed as for the CellROX Green assay above, except that 2 µM of probe was used and images acquired directly using Zen v3.4 after mounting the slides using a confocal rotary disk inverted microscope from Zeiss equipped with a Yokogawa CSU-1 module using a 20X objective. Some cells were treated with the GPX4 inhibitor RSL3 (0.3μM or 1μM) for 8 hours at 37 °C before loading the probe. The signal intensity and the ratio of the green signal (oxidized probe) to red signal (reduced probe) was measured and calculated using Fiji (version 2.17.0).

### Immunoprecipitation and Western blotting

Liver tissue pieces (∼50 mg) were homogenized in lysis buffer [20mM Tris-HCl (pH 7.5), 150mM NaCl, 1mM EDTA (pH 8.0), 1mM EGTA, 2.5mM NaPyrophosphate, 1mM β-glycerophosphate, 10mM NaF, 1% Triton, 1mM phenylmethylsulfonyl fluoride (PMSF) and protease inhibitors (4693132001; Roche Diagnostics)]. To immunoprecipitate Gla proteins, the protein extracts (0.2 or 1 mg) were incubated with 5µg of rabbit anti-Gla antibodies at 4°C overnight, as previously described ^14^ . Then, the extracts were incubated with Protein A-Agarose beads (11719408001; Roche Diagnostics) for 3 hours and washed 4 times with lysis buffer. Laemmli buffer was added to the immunoprecipitated proteins, heated at 70 °C for 10 minutes and loaded on a 7.5% polyacryramide Tris-Glycine gel. The standard western blot procedure was performed to detected proteins using appropriate primary and secondary antibodies. The antibodies used in this study are listed in Supplementary Table 5.

### RNA isolation and qPCR

RNA extraction and gene expression in liver was performed as we have done previously ^14^. Primers used in this study are listed in Supplementary Table 6.

### RNA-sequencing

Total RNA was extracted from ∼30 mg of liver tissue using TRIzol reagent (Thermo Fisher). Further purification of RNA and elimination of DNA were performed using Zymo-Spin IIICG columns according to the manufacturer’s instructions (ZymoResearch, Irvin, CA). Evaluation of RNA integrity was conducted using the 2100 Bioanalyzer system (Agilent Technologies) with the RNA 6000 Nano kit. Libraries were prepared from an equal quantity of total RNA (e.g., 1000ng or 900ng) for each sample. Poly(A)+ transcripts were enriched from 1 mg of total RNA using the NEBNext Poly(A) Magnetic isolation module (E7490; NEB), followed by library preparation using the KAPA stranded RNA-seq library preparation Hyperprep kit (KR0934; Roche Diagnostics) and the TruSeq DNA library Prep LT kit (Illumina), using respective manufacturer’s protocols. Library size distribution was assessed on a 2100 Bioanalyzer system (Agilent Technologies) and libraries were quantified by qPCR. Equimolar libraries were sequenced in pair-end-reads (PE50 or PE100), on a Novaseq 6000 system (Illumina), with a SPrime or S4 flowcell and a coverage of 50 million fragments per library. The library preparation was performed at Montreal Clinical Research Institute (IRCM) genomics platform, and the sequencing was carried out at the Génome Québec Innovation Center. The quality of the raw reads was assessed with FASTQC v0.11.8. After examining the quality of the raw reads, no trimming was deemed necessary. The reads were aligned to the GRCm38 genome with STAR v2.7.6a with more than 77% of reads uniquely mapped. Raw counts were computed using FeatureCounts v1.6.0 based on Ensembl mouse reference genome v101. Differential expression analysis was performed using the DESeq2 R package. Subsequent gene set enrichment and gene network analyses were performed using StringDB ^64^, and ShinyGO 0.80 ^65^, accessing Biological Process and Cellular Component (Gene Ontology). Enrichment was considered significant if the false discovery rate was <0.05, with only networks containing 800 or fewer genes were included in the analysis and top 10-20 pathways were shown. Previously published transcriptomic data from control subject and subjects with MASLD and MASH was used to investigate the overlap between the deregulated gene signature in human and *Vkorc1l1*-deficient mouse model ^20^. Given the very large number of gene deregulated in human MASLD and MASH, we focused the analysis on deregulated genes with adjusted P value of £ 0.000005. The overlap between the different transcriptomes was determined using jvenn (http://jvenn.toulouse.inra.fr/app/example.html). The statistical significance between each pair of comparisons was computed using an online tool (http://nemates.org/MA/progs/overlap_stats.html).

### Genetic association analyses

A meta-analysis of genome-wide association study (GWAS) of MASLD totalling 16,532 cases and 1,240,188 controls from Estonian Biobank, deCODE genetics, UK Biobank, INSPIRE+HerediGene (USA), FinnGen study v8, the EPoS Consortium, and eMERGE Network ^21,23,66^ was used to access the impact of *GUSB* and *VKORC1L1* gene expression on metabolic dysfunction-associated steatotic liver disease (MASLD) diagnosis. The GWAS of liver fat derived from abdominal magnetic resonance imaging (MRI) using deep learning (proton density fat fraction) was used as a validation of results obtained for MASLD. This GWAS includes data of 38,000 participants from the UK Biobank and the summary statistic are publicly available from the original study ^67^. eQTL summary statistics for the liver expression of *GUSB* and *VKORC1L1* were obtained as previously described by Gobeil et al. ^22^. All genetic analyses were performed using R version 4.3.3. We applied the Summary-data-based Mendelian Randomization (SMR) software v1.3.1 ^68^ to investigate whether the genetic association between metabolic dysfunction-associated steatotic liver disease (MASLD) and nearby SNPs is mediated through the expression of *GUSB* or *VKORC1L1*. SMR analyses were performed using default settings. To further evaluate pleiotropy versus LD, the HEIDI test was applied with default settings (≤ 20 SNPs in the cis region; window size 2 Mb). Associations with HEIDI p > 0.05 were considered consistent with mediation rather than confounding by LD. We applied PWCoCo (v1.1.0), a statistical framework that integrates conditional analyses (GCTA-COJO) with colocalization (coloc) to test whether the same causal variant underlies both a molecular trait and a complex trait. PWCoCo performs pairwise colocalization of conditionally independent signals using summary-level GWAS and QTL data, accounting for LD. Analyses of MASLD GWAS signals and liver eQTLs for *GUSB* and *VKORC1L1* were conducted within ±1 Mb of the lead eQTL, using LD estimates from the 1000 Genomes Project (European ancestry). Default priors were applied (p = 1×10, p = 1×10, p = 1×10), and colocalization was considered robust when the posterior probability for a shared causal variant (H) exceeded 0.8.

### Institutional review board approval

Patients of the Quebec Obesity Biobank provided informed consent to participate to this institutional biobank. The study was conducted in accordance with the Declaration of Helsinki and approved by the Institutional Review Board (Ethics Committee) of Institut universitaire de cardiologie et de pneumologie de Québec-Université Laval (IUCPQ-UL) (approval number 2021-3656; date of approval: June 17th 2021). UK Biobank received approval from the British National Health Service, North West - Haydock Research Ethics Committee (16/NW/0274). The analyses in the UK Biobank were performed using data application number 25205. All genotype and phenotype data were collected according to an informed consent obtained at the baseline assessment from all participants.

### Statistical analysis

Detailed statistical analyses for each experiment is provided in the corresponding figure legends. Data analysis was conducted using GraphPad Prism software (version 10.1.1). Results are expressed as mean ± standard error of the mean (SEM). For comparisons involving two independent groups, an unpaired two-tailed Student’s *t*-test was applied. Analyses involving multiple independent groups were evaluated using one-way ANOVA followed by Bonferroni’s post-tests. For experiments with repeated measurements, two-way ANOVA followed by Bonferroni’s multiple comparisons test was employed. Two-way ANOVA followed by Fisher’s least significant difference (LSD) post-test was performed for the comparison between multiple groups with two independent variables (e.g., ROS assay and in vivo VK_2_ treatment). Statistical significance is denoted in figures as follows: *p *<* 0.05*, \*\**p < 0.01, and *\*\*\**p < 0.001. All experiments were independently replicated a minimum of three times or included data from at least three separate animals of each genotype.

## Data and Resource Availability

The data sets generated and/or analyzed during the current study are available from the corresponding author upon reasonable request. The *Vkorc1^-/-^*, *Vkorc1l1^-/-^* and the *Vkorc1l1^flox/flox^* mouse lines and the mouse VKORC1 antibody used during the current study are available from the corresponding author upon reasonable request.

