## Supplementary material for "The vitamin K oxidoreductase VKORC1L1 prevents oxidative stress in hepatocytes and protects from MASLD and hepatocellular carcinoma": Extented data figures 1-5

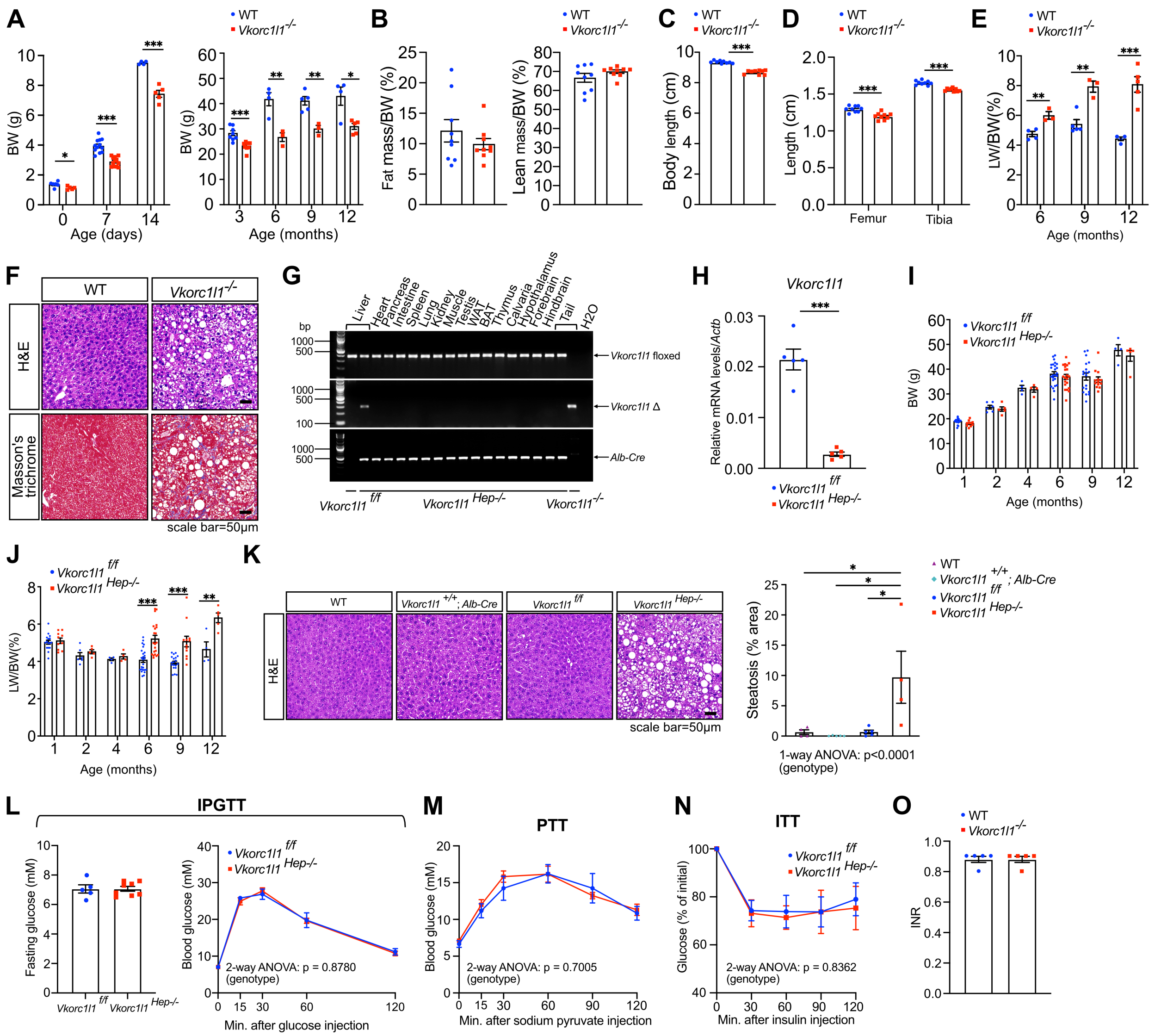


**Extended Data Fig 1: Metabolic phenotyping of *Vkorc1l1*$^{-/-}$ and *Vkorc1l1* $^{Hep-/-}$ mice. (A)** Body weight of wild-type (WT) and *Vkorc1l1*$^{-/-}$ male mice from P0 to 1 year of age (n = 3-12). **(B)** Fat and lean mass assessed by EchoMRI and normalized to body weight in 6-month-old WT and *Vkorc1l1*$^{-/-}$ male mice (n = 9). **(C)** Body length of 6-month-old WT and *Vkorc1l1*$^{-/-}$ male mice (n = 9). **(D)** Femur and tibia length of 6-month-old WT and *Vkorc1l1*$^{-/-}$ male mice (n = 9). **(E)** Liver weight normalized to body weight of 6-month-old WT and *Vkorc1l1*$^{-/-}$ male mice (n = 3-5). **(F)** H&E and Masson's trichrome staining of liver sections from 1-year-old WT and *Vkorc1l1*$^{-/-}$ male mice. **(G)** Genomic DNA from *Vkorc1l1*$^{f/f}$ and *Vkorc1l1* $^{Hep-/-}$ male mice was extracted, and PCR was used to detect the floxed and excised (Δ) *Vkorc1l1* alleles. **(H)** Gene expression analysis from liver tissue of 1-month old *Vkorc1l1*$^{f/f}$ and *Vkorc1l1* $^{Hep-/-}$ male mice using qPCR (n=5). **(I)** Body weight of 1- to 12-month-old *Vkorc1l1*$^{f/f}$ and *Vkorc1l1* $^{Hep-/-}$ male mice (n = 4-25). **(J)** Liver weight normalized to body weight of 1- to 12-month-old *Vkorc1l1*$^{f/f}$ and *Vkorc1l1* $^{Hep-/-}$ male mice (n = 4-25). **(K)** H&E staining of liver sections from 9-month-old WT, *Vkorc1l1* $^{+/+}$*; Alb-Cre, Vkorc1l1*$^{f/f}$ and *Vkorc1l1* $^{Hep-/-}$ male mice (n=4-17). Quantification represents % of steatosis over total liver area. **(L)** Intraperitoneal glucose tolerance test (IPGTT) performed on 4-month-old *Vkorc1l1*$^{f/f}$ and *Vkorc1l1* $^{Hep-/-}$ male mice (n = 5-8). **(M)** Pyruvate tolerance test (PTT) on 4-month-old *Vkorc1l1*$^{f/f}$

and *Vkorc1l1* $^{Hep-/-}$ male mice (n = 5-8). **(N)** Insulin tolerance test (ITT) in 4-month-old *Vkorc1l1*$^{f/f}$ and *Vkorc1l1* $^{Hep-/-}$ male mice (n = 7).-**(O)** Prothrombin time in 6-month-old WT and *Vkorc1l1*$^{-/-}$ male mice were measured using a Coagucheck and are shown as international normalized ratio (INR) (n = 5). Results represent the mean ± SEM. Unpaired, 2-tailed Student's t test was used in **(A-E, H-J, L and O)**. Ordinary one-way ANOVA with Bonferroni's post-tests was used in **(K)**. Two-way ANOVA with Bonferroni's post-tests was used in **(L-N)**. ***p < 0.001, **p < 0.01, *p < 0.05.

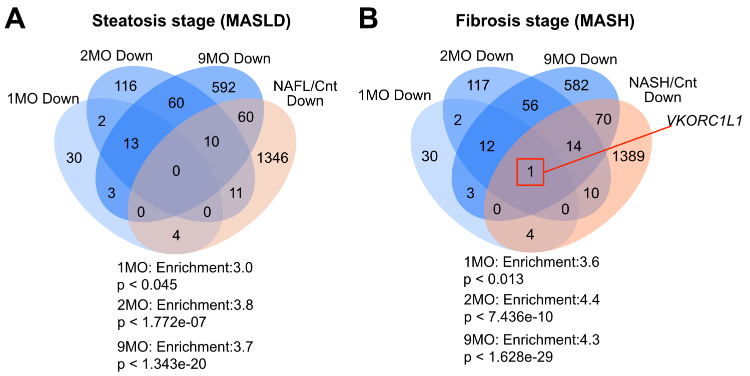


**Extended Data Fig 2: Overlap between dysregulated genes in *Vkorc1l1*-deficient livers and human MASLD/MASH gene signature.** Venn diagrams showing the overlap between down-regulated genes in 1-, 2- and 9-month-old *Vkorc1l1* deficient livers and down-regulated genes in human MASLD **(A)** and MASH **(B)**. Number of genes is shown for each overlap. *VKORC1L1* was found to be also significantly downregulated in human MASH.

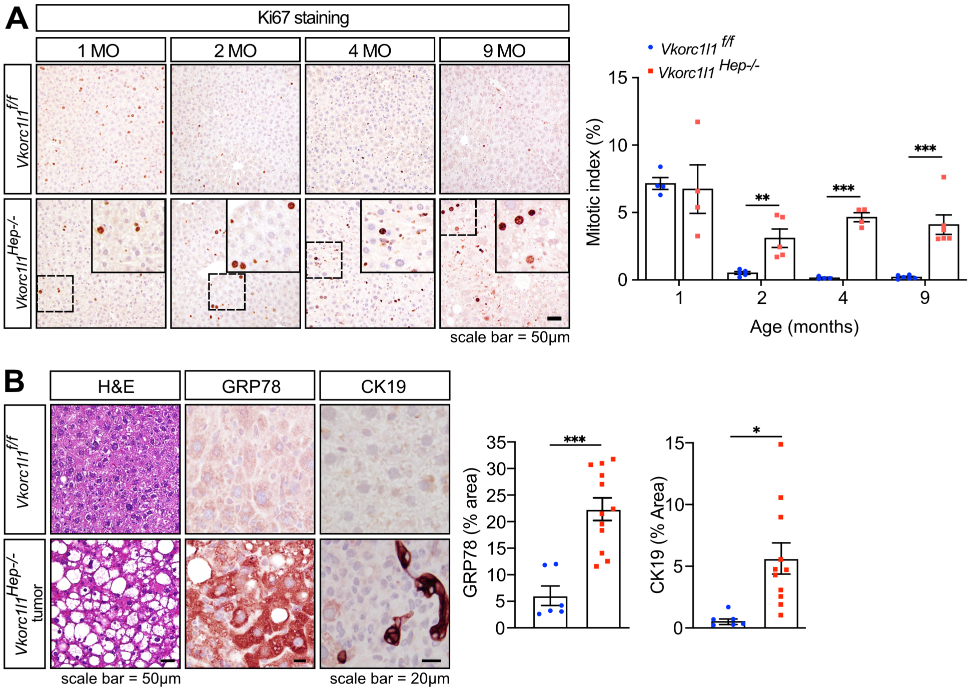


**Extended Data Fig 3: Characterization of proliferation and HCC in *Vkorc1l1*-deficient male mice. (A)** Ki67 immunohistochemistry on liver sections from *Vkorc1l1$^{f/f}$* and *Vkorc1l1* $^{Hep-/-}$ male mice at the indicated ages. The mitotic index was quantified as the percentage of Ki67+ cells over total cell number (n = 4-8). **(B)** H&E staining and GRP78 and CK19 immunohistochemistry of liver or tumor sections from 18- to 24-month-old *Vkorc1l1$^{f/f}$* and *Vkorc1l1* $^{Hep-/-}$ male mice. Quantifications are presented as the percentage of GRP78 or CK19 signal over total liver area. Results represent the mean ± SEM. Unpaired, 2-tailed Student's t test was used for the quantification in **(A)** and **(B)**. ***$p < 0.001$, **$p < 0.01$, *$p < 0.05$.

**A**

| Description | observed gene count | background gene count | strength | false discovery rate |
|---|---|---|---|---|
| Oxidoreductase activity | 22 | 787 | 0.61 | 0.00016 |
| Pheromone binding | 9 | 107 | 1.09 | 0.00025 |
| Oxidoreductase activity, acting on paired donors, with incorporation or reduction of molecular oxygen, reduced flavin or flavoprotein as one donor, and incorporation of one atom of oxygen | 8 | 80 | 1.16 | 0.00026 |
| Aromatase activity | 6 | 38 | 1.36 | 0.00042 |
| Oxidoreductase activity, acting on paired donors, with incorporation or reduction of molecular oxygen | 11 | 234 | 0.84 | 0.0007 |
| Steroid hydroxylase activity | 7 | 73 | 1.14 | 0.00077 |
| Monooxygenase activity | 9 | 159 | 0.92 | 0.0013 |
| Heme binding | 8 | 182 | 0.81 | 0.0202 |
| Oxidoreductase activity, acting on paired donors, with incorporation or reduction of molecular oxygen,NAD(P)H as one donor, and incorporation of one atom of oxygen | 5 | 65 | 1.05 | 0.041 |

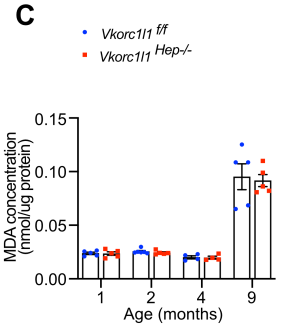


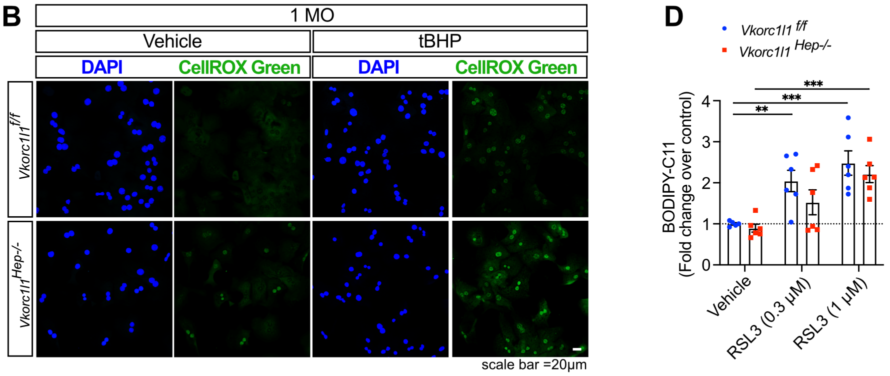


**Extended Data Fig 4: Impact of *Vkorc1l1* inactivation on oxidative stress, lipid peroxidation and ferroptosis. (A)** Gene Ontology (GO) analysis of dysregulated genes in 1-month-old *Vkorc1l1* $^{Hep-/-}$ male livers using molecular function (MF) terms. **(B)** Representative pictures of CellROX Green signal in primary hepatocytes isolated from 1-month-old *Vkorc1l1*$^{f/f}$ and *Vkorc1l1* $^{Hep-/-}$ male mice. **(C)** Lipid peroxidation was evaluated by measuring malondialdehyde (MDA) concentration in the liver from *Vkorc1l1*$^{f/f}$ and *Vkorc1l1* $^{Hep-/-}$ male mice at the indicated ages (n=4-5). **(D)** Lipid peroxidation was evaluated by using the ratiometric BODIPY$^{TM}$ 581/591 C11 fluorescent probe in primary hepatocytes isolated from 4-month-old *Vkorc1l1*$^{f/f}$ and *Vkorc1l1* $^{Hep-/-}$ male mice (n=5-6 experimental replicates per condition from 3 independent experiments). Quantifications represent the ratio of green (oxidized) over red (reduced) signal normalized to *Vkorc1l1*$^{f/f}$ hepatocytes treated with vehicle. Results represent the mean ± SEM. Multiple unpaired, 2-tailed Student's t test was used in **(C).** Two-way ANOVA with Bonferroni's post-tests was used in **(D)** ***p < 0.001, **p < 0.01, *p < 0.05.

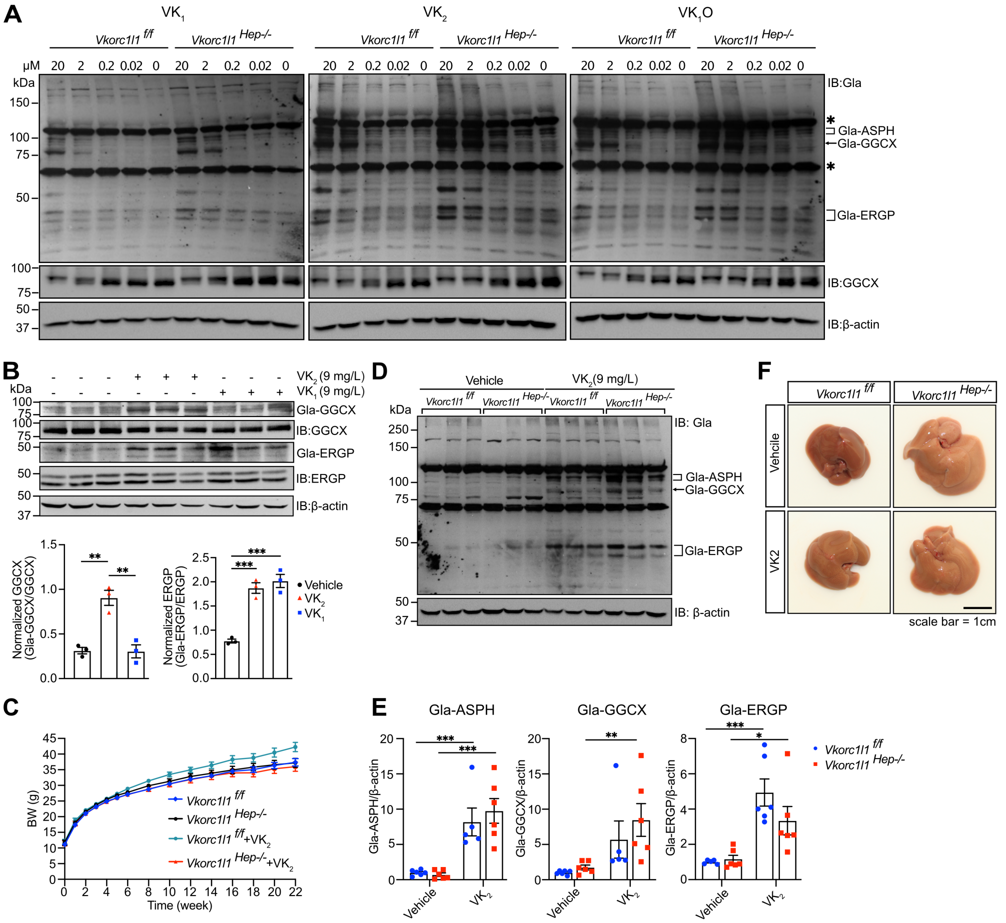


**Extended Data Fig 5: Effect of vitamin K supplementation on γ-carboxylation in hepatocytes and liver.** **(A)** Western blot analysis of γ-carboxylation in hepatocytes isolated from 4-month-old *Vkorc1l1$^{f/f}$* and *Vkorc1l1 $^{Hep-/-}$* male mice and treated with various concentration of phylloquinone (VK$_1$), menaquinone-4 (VK$_2$) or VK$_1$ 2,3-epoxide (VK$_1$O) for 24h using anti-Gla and anti-GGCX antibodies. β-actin was used as a loading control. **(B)** Western blot analysis of γ-carboxylation on liver extracts from 2-month-old C57BL/6J male mice treated with VK$_1$ or VK$_2$ in drinking water for 2 weeks using anti-Gla, anti-GGCX and anti-ERGP antibodies (n=3). β-actin was used as a loading control. **(C)** Body weight of *Vkorc1l1$^{f/f}$* and *Vkorc1l1 $^{Hep-/-}$* male mice during treatment with vehicle or VK$_2$ (n = 8-13). **(D)** Western blot analysis and quantification **(E)** of γ-carboxylation on liver extracts from 6-month-old *Vkorc1l1$^{f/f}$* and *Vkorc1l1 $^{Hep-/-}$* male mice treated with vehicle or VK$_2$ for 5 months (n = 5-6) using anti-Gla antibodies. β-actin was used as a loading control. **(F)** Gross morphology of liver from 6-month-old *Vkorc1l1$^{f/f}$* and *Vkorc1l1 $^{Hep-/-}$* male mice treated with vehicle or VK$_2$ for 5 months. Results represent the mean ± SEM. Ordinary one-way ANOVA with Bonferroni's post-tests was used in **(B)**. Two-way ANOVA with Bonferroni's post-tests was used in **(C).** Two-way ANOVA with Fisher's LSD was used in **(E)**. ***$p < 0.001$, **$p < 0.01$, *$p < 0.05$.
